# Activation of a signaling pathway by the physical translocation of a chromosome

**DOI:** 10.1101/2021.03.08.434431

**Authors:** Mathilde Guzzo, Allen G. Sanderlin, Lennice K. Castro, Michael T. Laub

## Abstract

In every organism, the cell cycle requires the execution of multiple cellular processes in a strictly defined order. However, the mechanisms used to ensure such order remain poorly understood, particularly in bacteria. Here, we show that the activation of the essential CtrA signaling pathway that triggers cell division in *Caulobacter crescentus* is intrinsically coupled to the successful initiation of DNA replication via the physical translocation of a newly-replicated chromosome, powered by the ParABS system. We demonstrate that ParA accumulation at the new cell pole during chromosome segregation recruits ChpT, an intermediate component of the CtrA signaling pathway. ChpT is normally restricted from accessing the selective PopZ polar microdomain until the new chromosome and ParA arrive. Consequently, any disruption to DNA replication initiation prevents the recruitment of ChpT and, in turn, cell division. Collectively, our findings reveal how major cell-cycle events are coordinated in *Caulobacter* and, importantly, how the physical translocation of a chromosome triggers an essential signaling pathway.

## Introduction

In all domains of life, cells must ensure that DNA replication, chromosome segregation, and cell division occur in the right order. A failure to coordinate these cell-cycle processes can lead to genome instability or cell death (Storchova and Pellman, 2004). In eukaryotes, numerous checkpoint systems ensure that each step of the cell cycle has completed before proceeding to the next one (Hartwell and Weinert, 1989; Murray, 1992). Few *bona fide* checkpoints have been identified in bacteria (Rudner and Losick, 2001) and the mechanisms that ensure orderly progression through the bacterial cell cycle remain largely unknown.

The *α*-proteobacterium *Caulobacter crescentus* has a tightly regulated cell cycle with DNA replication occurring only once per cell cycle under all growth conditions (Marczynski, 1999). *C. crescentus* also features an asymmetric cell division that generates two distinct daughter cells: a sessile stalked cell that immediately initiates replication and a motile swarmer cell that cannot initiate replication until it differentiates into a stalked cell (Fig. 1A). The initiation of DNA replication requires (i) the inactivation of CtrA, a response regulator that can directly silence the origin of replication (Domian *et al*., 1997; Quon *et al*., 1998), and (ii) accumulation of ATP-bound DnaA, the conserved replication initiator protein in bacteria (Gorbatyuk and Marczynski, 2001; Fernandez-Fernandez *et al*., 2011; Jonas *et al*., 2011). CtrA must then be produced *de novo* and activated by phosphorylation after DNA replication has initiated. In predivisional cells, CtrA directly promotes the expression of nearly 100 target genes critical to many cellular processes, including cell division (Laub *et al*., 2000, 2002). If DNA replication is blocked, CtrA is not produced (Wortinger *et al*., 2000) and cell division does not occur (Iniesta *et al*., 2010b), but the mechanism responsible for coupling replication to CtrA activation has not been identified.

**Figure 1:**
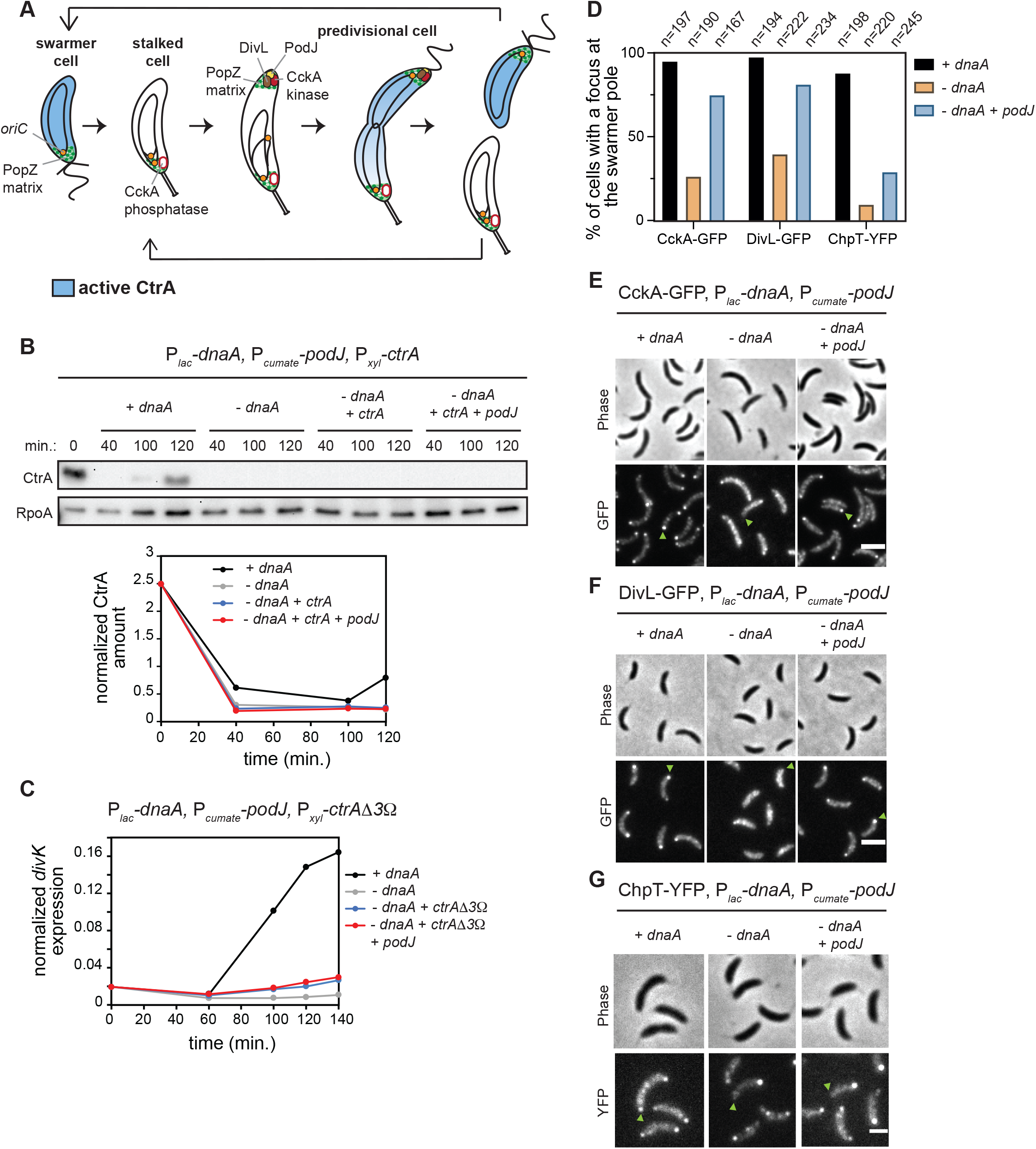
Polarization of CckA and DivL is not sufficient to activate CtrA if DNA replication is inhibited. (A) Schematic of the *Caulobacter crescentus* cell cycle. (B) CtrA immunoblots in synchronized cells expressing *dnaA* (+IPTG) or depleted of *dnaA* (-IPTG) with ectopic expression of *ctrA* (+0.075% xyl) and *podJ* (+cumate) when indicated. Times indicate minutes post-synchronization. Graph shows CtrA band intensity normalized to RpoA control. (C) mRNA levels of the CtrA-activated gene *divK* measured by qRT-PCR and normalized to *rpoA* mRNA levels in cells expressing *dnaA* (+IPTG) or depleted of *dnaA* (-IPTG) with ectopic expression of the proteolytically stable mutant *ctrAΔ3Ω* (+0.075% xyl) and *podJ* (+cumate) when indicated. (D-G) Localization of CckA-GFP (E), DivL-GFP (F) and ChpT-YFP (G) in fixed predivisional cells (90 min. post-synchronization) expressing *dnaA* (+IPTG) or depleted of *dnaA* (-IPTG) with or without ectopic expression of *podJ* (+cumate). Green arrows indicate the swarmer pole of one representative cell for each condition. (D) Histogram represents the percentage of cells with detectable foci of CckA-GFP, DivL-GFP, or ChpT-YFP at the swarmer pole (includes bipolar and unipolar clusters at the swarmer pole) in each condition. Total number of cells examined from two biological replicates is indicated in each case. Scale bar = 2 μm.

CtrA activity is controlled by a phosphorelay from the histidine kinase CckA to the histidine phosphotransferase ChpT to CtrA (Biondi *et al*., 2006) (Fig. S1A). Following DNA replication, CckA is recruited to the nascent swarmer pole of predivisional cells where its kinase activity is stimulated by the atypical histidine kinase DivL (Jacobs *et al*., 1999; Chen *et al*., 2009; Angelastro *et al*., 2010; Iniesta *et al*., 2010a; Tsokos *et al*., 2011) (Fig. 1A, S1A). CckA and ChpT also drive phosphorylation of CpdR to prevent it from promoting CtrA degradation (Jenal and Fuchs, 1998; Biondi *et al*., 2006; Iniesta *et al*., 2006; Chien *et al*., 2007) (Fig. S1A). The localization of CckA and DivL to the swarmer pole and CckA kinase activity depend on DNA replication initiation (Iniesta *et al*., 2010b), but the underlying mechanism is not known. Additionally, whether the localization of CckA and DivL is the key means of coupling DNA replication to CtrA activation and subsequent cell division has not been established.

Here, we show that swarmer pole localization of CckA and DivL can be restored in the absence of DNA replication by ectopically producing the scaffold protein PodJ. Surprisingly though, this polarization of CckA and DivL is not sufficient to activate CtrA. Instead, we find that the activation of CtrA after DNA replication has initiated depends on the recruitment of the intermediate component of the phosphorelay, ChpT, to the swarmer pole by the chromosome segregation machinery. Replication initiation occurs near the stalked pole in *Caulobacter*. One of the duplicated chromosomes is then translocated to the opposite cell pole via a complex interplay between the nucleoid-bound ATPase ParA and the protein ParB bound to origin-proximal *parS* sites (Toro *et al*., 2008; Lim *et al*., 2014). We show that the accumulation of nucleoid-free ParA at the swarmer pole once chromosome translocation has completed drives the recruitment of ChpT, thereby completing the phosphorelay from CckA to CtrA. When DNA replication initiation is blocked or chromosome translocation is disrupted, ChpT fails to accumulate at the swarmer pole, preventing cell division. Thus, our work now reveals the molecular basis of a substrate-product relationship, rather than a checkpoint, by which replication and cell division are coupled in *Caulobacter crescentus*. More broadly, our results also demonstrate how a physical cue, in this case the translocation of a chromosome, can serve as the activating signal for a complex signal transduction pathway.

## Results

### CtrA activation in predivisional cells depends on DNA replication

Prior studies have indicated that CtrA activation in predivisional cells depends on DNA replication initiation (Wortinger *et al*., 2000; Iniesta *et al*., 2010b). To corroborate this observation and establish a genetic system for further studying it, we used a strain in which the endogenous *dnaA* promoter was replaced by the *E. coli lac* promoter and the *lacI* gene was integrated at the *hfa* locus (Badrinarayanan *et al*., 2015). We grew cells in the absence of IPTG for 90 minutes to deplete DnaA, and then synchronized and released cells into fresh minimal medium (M2G+) with or without IPTG to induce *dnaA* expression or not, respectively. In the presence of DnaA, DNA replication initiated within 40 min., as judged by flow cytometry (Fig. S1B), whereas in DnaA-depleted cells, DNA replication had not initiated even by the end of the time-course (120 min.) (Gorbatyuk and Marczynski, 2001) (Fig. S1B).

To test whether CtrA was activated in the absence of DNA replication, we first used Western blots to measure CtrA abundance. The accumulation of CtrA in predivisional cells depends on a positive feedback loop wherein active, phosphorylated CtrA promotes its own transcription and accumulation (Quon *et al*., 1998). In cells producing DnaA, CtrA accumulated in late predivisional cells, ∼120 min. post-synchronization, as expected (Fig. 1B), but in cells lacking DnaA, CtrA did not accumulate (Fig. 1B) because *ctrA* is not transcribed in cells that lack DnaA (Holtzendorff *et al*., 2004). Therefore, we ectopically expressed *ctrA* from a xylose-inducible promoter on a plasmid in cells also depleted of DnaA, but CtrA still did not accumulate by 120 min. post-synchronization in the presence of xylose, presumably because it gets degraded (Fig. 1B).

The lack of CtrA accumulation in cells depleted of DnaA supports the notion that CtrA activation is dependent on DNA replication initiation. However, cells lacking DnaA could be competent for CtrA phosphorylation, but also be actively degrading CtrA. To rule out this possibility, we expressed a constitutively stable variant, *ctrAΔ3Ω* that can still be regulated by phosphorylation (Domian *et al*., 1997) (Fig. S1C), in cells depleted of DnaA. We now observed an accumulation of this stabilized CtrA (Fig. S1D). To assess CtrA phosphorylation, we used qRT-PCR to measure mRNA levels of the CtrA target gene *divK*. In cells expressing *dnaA, divK* mRNA levels increased substantially in predivisional cells (Fig. 1C). In contrast, for cells depleted of DnaA, *divK* expression remained low throughout the cell cycle either with or without CtrAΔ3Ω being produced (Fig. 1C). Thus, even if abundant in cells, CtrA is not phosphorylated in the absence of DnaA and DNA replication initiation.

### Known regulators cannot account for a failure to activate CtrA after inhibiting DNA replication

DnaA promotes the transcription of *gcrA* (Hottes *et al*., 2005), a key transcriptional regulator in *Caulobacter* (Haakonsen *et al*., 2015). Thus, in the absence of DnaA, either GcrA or an unknown GcrA-regulated factor that is required for CtrA activation could be missing. To test this possibility, we ectopically expressed *gcrA-3xflag* (Fig. S1E) and *ctrA* in cells depleted of DnaA. However, CtrA did not accumulate in predivisional cells of this strain (Fig. S1E), and *divK* was not activated even when expressing non-degradable CtrAΔ3Ω (Fig. S1F). These results indicate that a lack of GcrA in the absence of DnaA is not responsible for blocking CtrA activation.

Next, we considered whether inhibiting DNA replication initiation somehow inactivated CckA, which can be directly inhibited by c-di-GMP (Lori *et al*., 2015). To test this model, we reduced c-di-GMP levels in cells depleted of DnaA by expressing a phosphodiesterase (PA5295) from *Pseudomonas aeruginosa* (Abel *et al*., 2013). In this strain, low c-di-GMP levels prevented CtrA degradation at the swarmer-to-stalk cell transition, so CtrA remained present throughout the cell cycle (Fig. S1G). However, *divK* expression was not upregulated in the absence of DnaA when expressing this phosphodiesterase (Fig. S1H). Thus, we infer that blocking DNA replication does not prevent CtrA activation through c-di-GMP-dependent inhibition of CckA.

Finally, we tested whether SciP, a protein that inhibits CtrA transcriptional activity in swarmer cells (Gora *et al*., 2010), somehow accumulated following a block to replication initiation. However, SciP was still properly degraded 30 min. after synchronization and remained low in cells depleted of *dnaA* (Fig. S1I), indicating that an accumulation of SciP does not explain the inhibition of CtrA if DNA replication fails to initiate.

### Polar localization of DivL and CckA is not sufficient to activate CtrA following a block to DNA replication

CtrA phosphorylation in predivisional cells depends on the polar localization of the histidine kinases CckA and DivL (Jacobs *et al*., 1999; Chen *et al*., 2009; Angelastro *et al*., 2010; Iniesta *et al*., 2010a; Tsokos *et al*., 2011). This localization of CckA and DivL depends on DNA replication (Iniesta *et al*., 2010b), and we confirmed that in the absence of DnaA, CckA-GFP and DivL-GFP do not localize to the nascent swarmer pole of predivisional cells (Fig. 1D-F). We then wanted to test if this defect in CckA and DivL localization was the limiting step for CtrA activation in the absence of DnaA. Notably, DnaA directly upregulates *podJ* expression after replication initiation (Hottes *et al*., 2005) (Fig. S1C) and the polar localization of DivL and CckA both depend on PodJ (Curtis *et al*., 2012). Thus, to test if PodJ was the limiting factor for CckA and DivL localization (and ultimately for CtrA activation) in the absence of DnaA, we ectopically expressed *podJ* using a cumate-inducible promoter (Kaczmarczyk *et al*., 2013) in the *dnaA* depletion strain (Fig. S1C). Inducing PodJ was sufficient to restore CckA-GFP and DivL-GFP to the nascent swarmer pole in the absence of DnaA (Fig. 1D-F; -*dnaA* +*podJ* condition). To test if CtrA activation was restored, we took cells depleted of DnaA and ectopically expressed *ctrA* and *podJ*. CtrA levels remained low even 120 min. after synchronization (Fig. 1B) and *divK* expression was not upregulated, even when expressing the *ctrAΔ3Ω* stable variant (Fig. 1C). These results indicate that PodJ production is sufficient to polarize CckA and DivL in the absence of DNA replication, but that CtrA is still not phosphorylated.

Importantly, the phosphorylation of CtrA requires autophosphorylated CckA to transfer a phosphoryl group to the histidine phosphotransferase ChpT which then serves as the direct phosphodonor to CtrA (Biondi *et al*., 2006) (Fig. S1A). ChpT was not considered in previous studies of how DNA replication initiation affects CtrA. To test whether ChpT localization depends on replication initiation, we engineered the DnaA-depletion strain to produce ChpT-YFP from its native genomic locus. In the presence of DnaA, ChpT-YFP foci were seen at the nascent swarmer pole of predivisional cells (Fig. 1D,G). In contrast, for cells depleted of DnaA, no foci were seen at that pole, only the stalked pole (Fig. 1D,G). Thus, like CckA and DivL, ChpT localization to the nascent swarmer pole depends on the successful initiation of DNA replication. However, in sharp contrast to our observations for CckA and DivL, the ectopic production of PodJ was not sufficient to localize ChpT-YFP (Fig. 1D,G). Thus, ChpT localization to the swarmer pole is not determined simply by the localization of CckA. Additionally, this finding indicates that localization of ChpT may be the limiting step in coupling CtrA activation to DNA replication initiation.

### Only partial replication of the chromosome is required for CtrA activation

The findings presented thus far suggest that the requirement for DnaA in localizing ChpT to the swarmer pole, and in turn activating CtrA, is not related to its role as a transcription factor. Thus, we favored the possibility that something about the act of DNA replication itself, which is triggered by DnaA, is required to activate CtrA. To further explore how replication controls CtrA activation, we examined a strain in which wild-type DnaA was depleted but ectopically produces DnaA(R357A), which is likely locked in the active, ATP-bound form (Fernandez-Fernandez *et al*., 2011; Jonas *et al*., 2011). This strain overinitiates DNA replication and the chromosome content per cell exceeds 2N, as judged by flow cytometry (Fig. S2A). Using qPCR at seven loci along the chromosome (Fig. 2A), we found that cells producing DnaA(R357A), accumulated up to 5 copies of genomic regions near the origin (Fig. 2A). In contrast, cells producing wild-type DnaA had at most 2 copies, as expected for wild-type *Caulobacter* (Marczynski, 1999) (Fig. 2A). Despite the high initiation rate of cells producing DnaA(R357A), the genomic copy number was less than 2 at 0.67 Mb and unreplicated beyond 1 Mb (Fig. 2A). Thus, this strain enabled us to ask whether only partial (∼1/3) replication of the chromosome was sufficient to activate CtrA.

**Figure 2:**
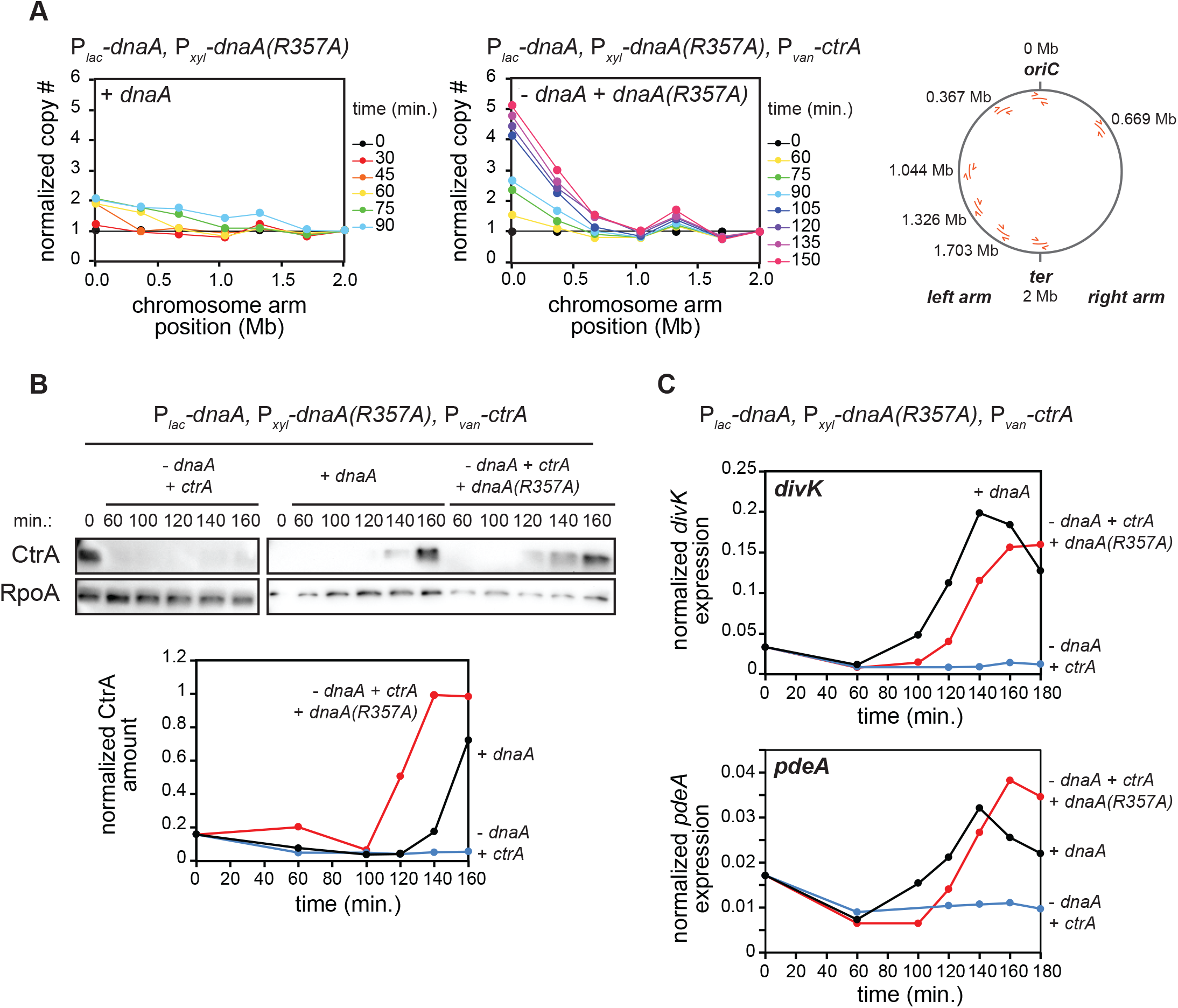
Replication of the full chromosome is not required for CtrA activation. (A) qPCR on genomic DNA extracted from cells expressing *dnaA* (+IPTG) or depleted of *dnaA* and expressing the *dnaA(R357A)* ATP-locked mutant (-IPTG +xyl) at different timepoints post-synchronization. Map of *C. crescentus* chromosome (right) shows the localization of primers used to assess replication and their distances from *oriC*. Samples were normalized internally to DNA levels close to the terminus (*CCNA_01869*) and each timepoint was then normalized to t=0 min. to evaluate fold-change from 1N. Note that for the ‘+*dnaA’* condition, samples after t=90 min. were excluded because DNA close to the terminus has been replicated. (B) CtrA protein levels at the times indicated post-synchronization in cells expressing *dnaA* (+IPTG) or depleted of *dnaA* (-IPTG) with ectopic expression of wild-type *ctrA* (+van) and with (+xyl) or without ectopic expression of *dnaA(R357A)*. Graph shows CtrA protein band intensity normalized to RpoA. (C) mRNA levels of the CtrA-activated genes *divK* and *pdeA* measured by qRT-PCR and normalized to *rpoA* mRNA levels for cells treated as in (B).

We first examined *ctrA* transcription, finding that it remained relatively low in cells producing DnaA(R357A) (Fig. S2B) despite the accumulation of GcrA (Fig. S2C), possibly because the *ctrA* locus at 0.764 Mb remains fully methylated, which reduces its transcription (Holtzendorff *et al*., 2004). We therefore placed *ctrA* under the control of a vanillate-inducible promoter in cells also engineered to deplete wild-type DnaA and produce DnaA(R357A). We depleted DnaA for 90 minutes, synchronized cells, and released them into a medium that enables repression of wild-type DnaA and induction of DnaA(R357A) and *ctrA* (Fig. S2D). CtrA now strongly accumulated in late predivisional cells despite the absence of full chromosome replication (Fig. 2B). Additionally, the CtrA-regulated genes *divK* and *pdeA* were strongly upregulated in cells expressing both *dnaA(R357A)* and *ctrA* compared to cells depleted of DnaA (Fig. 2C). We conclude that replication of the entire chromosome is not required for CtrA activation. Either the act of replication initiation itself or an event coupled to the earliest stages of replication is required.

### Disrupting chromosome segregation prevents CtrA activation

Our experiments with the overinitiating *dnaA(R357A)* strain suggested that replication of the first third of the chromosome is sufficient to trigger CtrA activation in predivisional cells (Fig. 2). Notably, chromosome segregation is initiated upon replication of the *parS* site, which is located just 8 kb-away from *oriC*, the chromosomal origin of replication (Toro *et al*., 2008). After the *parS* locus is duplicated, one of the copies separates away from the stalked pole, and then ParA and ParB drive its rapid translocation across the cell to the nascent swarmer pole (Ptacin *et al*., 2010; Schofield *et al*., 2010; Shebelut *et al*., 2010; Lim *et al*., 2014) (Fig. 3A). Thus, we decided to investigate whether segregation of the chromosome is a critical step in CtrA activation. Such a role for chromosome segregation was previously dismissed (Iniesta *et al*., 2010b) because synthesizing ParA(K20R), a variant of ParA that blocks chromosome segregation without altering replication (Toro *et al*., 2008) (Fig. S3A), did not affect CckA-GFP localization. However, the impact of ParA(K20R) on CtrA activation and ChpT localization was not tested.

**Figure 3:**
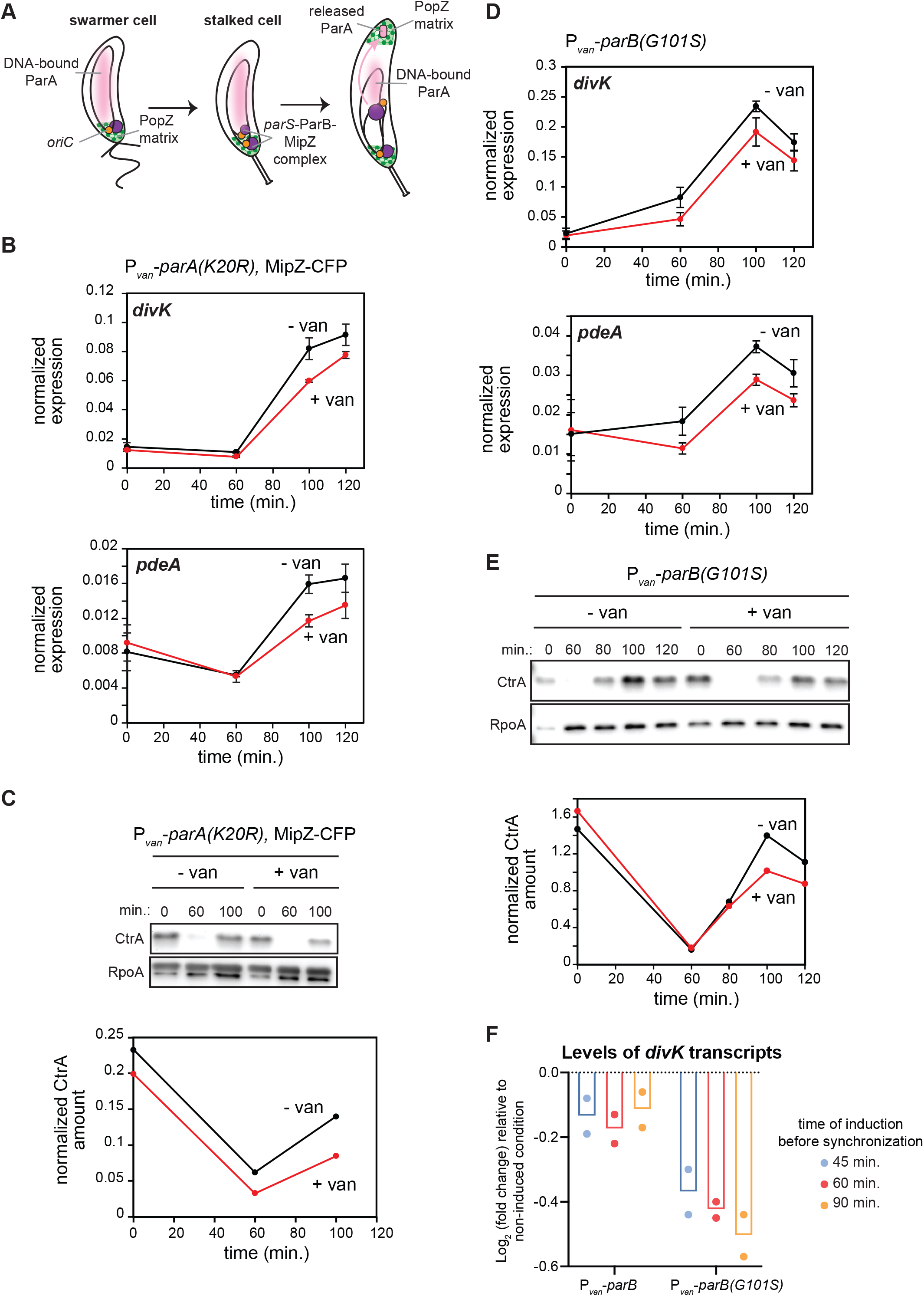
CtrA activation in predivisional cells is reduced when chromosome segregation is perturbed. (A) Schematic of chromosome segregation during the *C. crescentus* cell cycle. (B) mRNA levels of the CtrA-activated genes *divK* and *pdeA* measured by qRT-PCR and normalized to *rpoA* mRNA levels at the times indicated post-synchronization in cells expressing (+van) or not (-van) parA*(K20R)*. Expression of *parA(K20R)* was induced 60 min. pre-synchronization and post-synchronization with 500 μM vanillate. Data represent the mean ±SD of three biological replicates. (C) CtrA protein levels at the times indicated post-synchronization in cells from the same conditions as in (B). Graph shows CtrA protein band intensity normalized to RpoA. (D) mRNA levels of the CtrA-activated genes *divK* and *pdeA* measured by qRT-PCR and normalized to *rpoA* mRNA levels at the times indicated post-synchronization in cells expressing (+van) or not (-van) the spreading-deficient *parB(G101S)* mutant. Expression of *parB(G101S)* was induced 60 min. pre-synchronization and post-synchronization with 500 μM vanillate. Data represent the mean ± SD of three biological replicates. (E) CtrA protein levels at the times indicated post-synchronization in cells from the same conditions as in (D). Graph shows CtrA protein band intensity normalized to RpoA. (F) Relative fold-change in mRNA levels of the CtrA-regulated gene *divK* measured by qRT-PCR and normalized to *rpoA* mRNA levels at timepoint 100 min. post-synchronization, when inducing *parB* or *parB(G101S)* 45 min., 60 min. or 90 min. before synchronization compared to the uninduced condition. Shown are averages from two biological replicates with the individual datapoints for each replicate.

We first tested if disrupting chromosome segregation by ectopically expressing *parA(K20R)* would affect CtrA activation. To try and maximize the disruption of chromosome segregation but retain synchronizability of cells, we induced *parA(K20R)* 60 minutes before synchronizing cells (Fig. 3B-C). We found that the expression levels of two CtrA-activated genes, *divK* and *pdeA,* and CtrA protein levels were both modestly, but significantly reduced in predivisional cells (100 min. post-synchronization) expressing *parA(K20R)* compared to the non-induced condition (Fig. 3B-C). These results indicated that improper segregation of the chromosome impacts CtrA activation. ParA(K20R) did not completely eliminate CtrA activation, likely due to the heterogeneity of ParA(K20R) accumulation and an incomplete disruption of chromosome segregation, which we return to later.

To corroborate the results with ParA(K20R), we sought to perturb chromosome segregation, again without disrupting DNA replication, in an alternative way. To do this, we overexpressed the spreading-deficient mutant *parB(G101S)* (Tran *et al*., 2018), which contains a point mutation in the arginine-rich patch important for CTP binding (Jalal *et al*., 2020). Inducing *parB(G101S)* does not affect DNA replication (Fig. S3B), but prevents chromosome segregation, as indicated by a failure to segregate the ParB-associated protein MipZ to the nascent swarmer pole (Fig. S3C). As with *parA(K20R),* inducing the expression of *parB(G101S)* 60 minutes before synchronization also modestly, but significantly, reduced the expression levels of two CtrA-regulated genes, *divK* and *pdeA,* and CtrA protein levels (Fig. 3D-E). Additionally, we observed that inducing *parB(G101S)* for increasingly longer times prior to synchronization (45 min., 60 min. or 90 min.) enhanced the effect on CtrA activation (Fig. 3F), suggesting that the reduction in CtrA activation is proportional to the accumulation of *parB(G101S)*. Notably, we did not observe this dose-response effect when expressing wild-type *parB* instead of *parB(G101S)* (Fig. 3F). Taken together, our results suggest that chromosome segregation is critical in triggering the proper accumulation of phosphorylated, active CtrA following DNA replication initiation.

### Disrupting chromosome segregation affects the localization of ChpT, but not CckA and DivL

To investigate the connection between chromosome translocation and the polarization of DivL, CckA, and ChpT, we induced *parA(K20R)* from a vanillate-inducible promoter in cells also expressing MipZ-CFP, CckA-GFP, DivL-GFP, or ChpT-sfGFP at their native genomic loci. When inducing *parA(K20R)* for 30 min. before synchronization, some cells can divide while some cannot and instead become elongated. The percentage of elongated cells when expressing *parA(K20R)* was 38% for MipZ-CFP (n=309), 50% for CckA-GFP (n=335), 36% for DivL-GFP (n=207) and 35% for ChpT-sfGFP (n=310). This heterogeneity likely reflects heterogeneity in ParA(K20R) accumulation, with elongated, undivided cells representing those with enough ParA(K20R) to disrupt chromosome segregation. We therefore focused on elongated cells that had not divided by 270 min. after synchronization in the following time-lapse experiments; for reference, non-induced cells handled identically divide after ∼150 min. (Fig. 4A). In the elongated cells, MipZ-CFP (which binds and labels ParB-*parS* complexes (Thanbichler and Shapiro, 2006)) localization to the new pole was strongly reduced, as expected if chromosome segregation is disrupted (Toro *et al*., 2008) (Fig. 4B-C). For both dividing and elongating cells, CckA-GFP and DivL-GFP accumulated at the new swarmer pole in almost all cells of the population (Fig. 4B, D, S4A) (Iniesta *et al*., 2010b), consistent with our findings that CckA and DivL localization depends primarily on PodJ accumulation at the pole (Fig. 1D-F), not DNA replication or subsequent chromosome segregation. In sharp contrast to CckA and DivL, only 30% of elongated cells producing ParA(K20R) had detectable ChpT-sfGFP foci at the nascent swarmer pole (Fig. 4B, E). These results suggest that, unlike CckA and DivL, ChpT localization to the swarmer pole depends on proper chromosome segregation.

**Figure 4:**
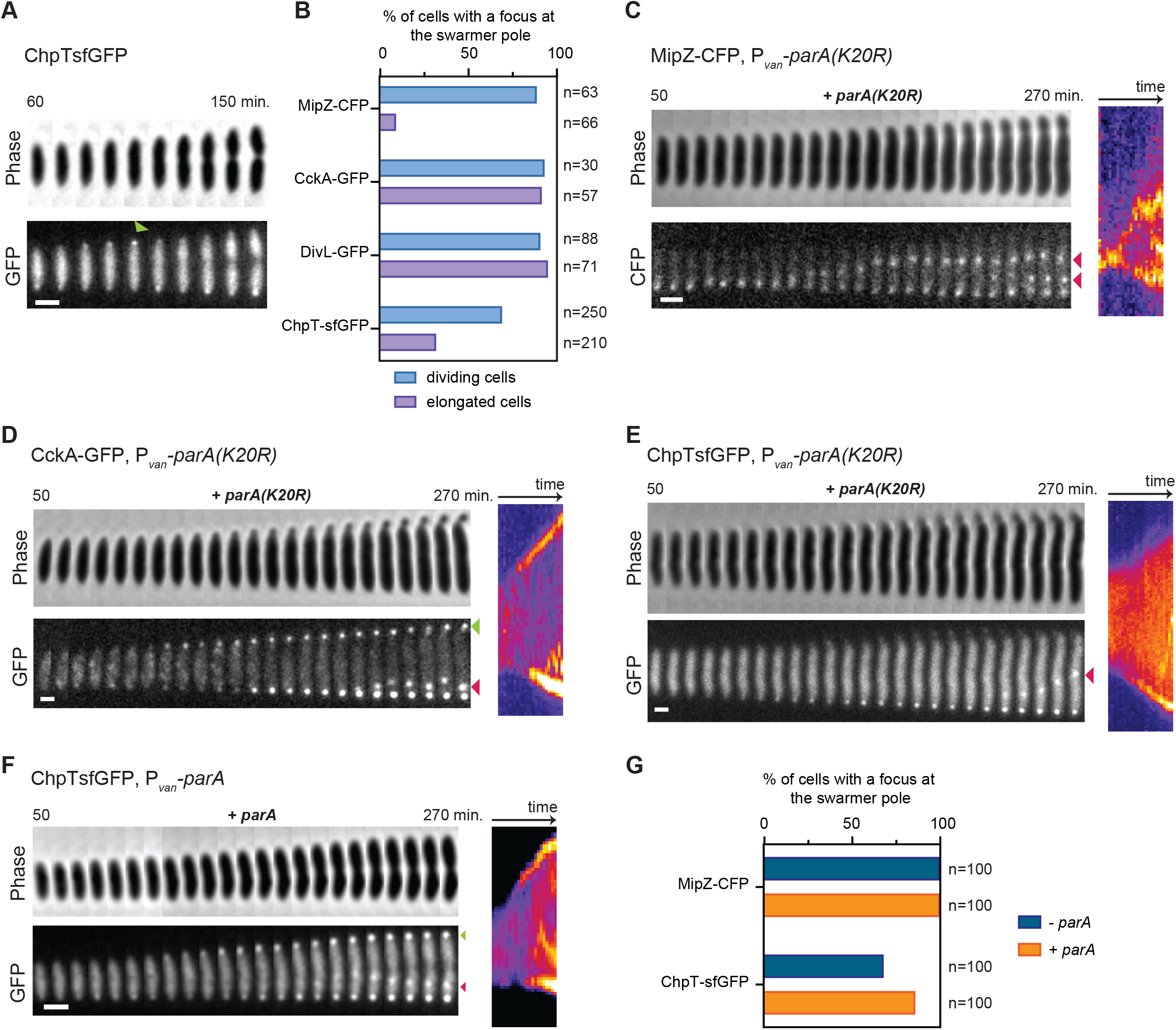
ChpT access to the new swarmer pole is blocked when chromosome segregation is disrupted. (A) Time lapse of ChpT-sfGFP dynamics in a wild-type cell, imaged at 10 min. intervals from 60 to 150 min. (before division) post-synchronization, with consecutive frames pasted side-by-side to generate the concatenated figure. Green arrow indicates ChpT-sfGFP transient localization at the new pole. Scale bar = 1 μm. (B) Percentage of cells with detectable foci at the swarmer pole at any time during time-lapse imaging of MipZ-CFP, CckA-GFP, DivL-GFP, and ChpT-sfGFP without *parA(K20R)* induction (dividing cells; -van) and with *parA(K20R)* induction (elongated cells only; +van). Expression of *parA(K20R)* was induced 30 min. before synchronization and after synchronization with 500 μM vanillate. Total number of cells examined from two biological replicates is indicated in each case. (C-E) Examples of a single elongated cell from each condition quantified in (B). Scale bars = 1 μm. (C) Time-lapse of MipZ-CFP dynamics within a single cell expressing the dominant negative *parA(K20R)* mutant, imaged at 10 min. intervals from 50 to 270 min. post-synchronization, with consecutive timeframes pasted side-by-side to generate the concatenated figure (left) and corresponding kymograph (right). Red arrows indicate MipZ-CFP internal clusters next to the newly replicated chromosomal origins. (D) Same as (C) but for CckA-GFP. Green arrow shows CckA-GFP localization at the new swarmer pole upon expression of *parA(K20R).* Red arrow shows CckA-GFP accumulation in a single internal cluster. (E) Same as (C) but for ChpT-sfGFP. Red arrow shows ChpT-sfGFP accumulation in a single internal cluster. (F) Timelapse of ChpT-sfGFP dynamics within a single cell expressing ectopic wild-type *parA*, imaged at 10 min. intervals from 50 to 270 min. post-synchronization, with consecutive timeframes pasted side-by-side to generate the concatenated figure (left) and corresponding kymograph (right). Expression of *parA* was induced 30 min. before synchronization and during imaging post-synchronization in the agarose pad with addition of 500 μM vanillate. Green arrow shows ChpT-sfGFP localization at the new swarmer pole upon expression of *parA.* Red arrow shows ChpT-sfGFP accumulation in a single internal cluster. Scale bar = 1 μm. (G) Percentage of cells with detectable foci at the swarmer pole at any time during time lapse imaging of ChpT-sfGFP (see Fig. 4F) and MipZ-CFP (see Fig. S4B) (n=100 cells for each condition) without (-van condition) or with *parA* induction (+van).

We also examined ChpT-sfGFP localization in cells overexpressing a wild-type copy of *parA* (Fig. 4F-G), which does not prevent chromosome translocation as with *parA(K20R)* (Fig. 4G, S4B). In these cells, ChpT-sfGFP now accumulated at the nascent swarmer pole (Fig. 4F-G). In fact, whereas ChpT-sfGFP localization to that pole is typically transient in WT cells (Fig. 4A) and not always easily detectable (Fig. 4B), overexpression of wild-type *parA* strongly increased the stability and detectability of ChpT-sfGFP at the new pole in most of the cells in the population (Fig. 4F-G), and resembled MipZ-CFP localization under the same conditions (compare Fig. 4F and S4B). Collectively, our results support the conclusion that localization of ChpT to the swarmer pole, and subsequent activation of CtrA, depends on successful chromosome translocation, which occurs shortly after replication initiation.

### ChpT localization to the new pole and access to the PopZ microdomain depends on ParA

For ChpT-sfGFP to localize to the swarmer pole, it must access the microdomain formed by the polar organizer protein PopZ, which excludes most cytoplasmic proteins (Ebersbach *et al*., 2008; Bowman *et al*., 2010) unless they bind a component of the microdomain (Lasker *et al*., 2020). The formation of the PopZ microdomain was not affected by the overexpression of *parA(K20R)* (Laloux and Jacobs-Wagner, 2013) (Fig. S4C), and ChpT-sfGFP appeared confined to the rest of the cell (Fig. 4E) suggesting that ChpT is excluded from the PopZ microdomain in the absence of chromosome translocation. Overexpressing *popZ* can expand the PopZ microdomain, especially at the old cell pole (Ebersbach *et al*., 2008; Bowman *et al*., 2010; Lasker *et al*., 2020). We induced the expression of *popZ* after synchronization (of cells also producing ParA(K20R) to disrupt chromosome segregation) and asked how this affected CckA and ChpT localization within that first cell cycle. CckA-GFP remained bipolar (Fig. 5A, S5A), suggesting that an excess of PopZ does not disrupt recruitment or retention of CckA in these conditions. In contrast, ChpT-sfGFP completely failed to accumulate at the swarmer pole following *popZ* overexpression, with a significant drop in the ChpT-sfGFP cytoplasmic signal and concomitantly stronger signal at the old, stalked pole (Fig. 5B, S5B). These results indicated that ChpT is indeed excluded from the swarmer pole microdomain and that an additional component is necessary for ChpT to access that pole. Additionally, overexpressing *popZ* has recently been shown to reduce CtrA activation (Lasker *et al*., 2020), consistent with a reduction of ChpT access to the new pole where phosphotranfer from CckA to ChpT to CtrA can occur.

**Figure 5:**
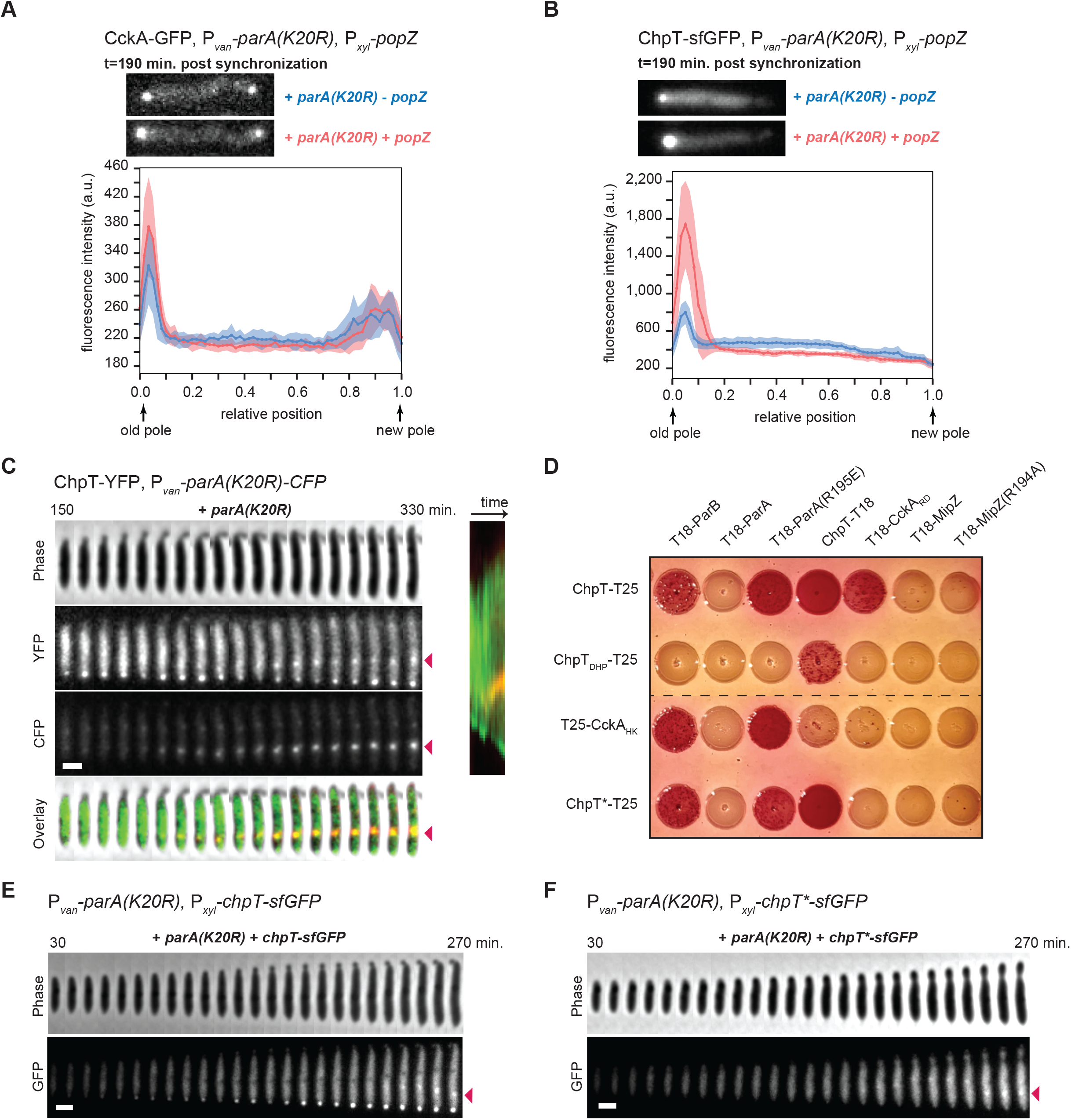
ParA not bound to DNA recruits ChpT to the swarmer pole. (A-B) Fluorescence micrographs (top) of CckA-GFP (A) or ChpT-sfGFP (B) in cells expressing *parA(K20R)* (+van) with (+xyl) or without (-xyl) induction of *popZ* post-synchronization (for full time lapses see Fig. S5A-B for CckA-GFP and ChpT-sfGFP, respectively). Graphs (bottom) show the average fluorescence intensities (solid line) from the old pole to the new pole from 15 cells for each condition. Shading around the solid line represents the SD. Scale bars = 1 μm. (C) ChpT-YFP internal cluster colocalizes with ectopically produced ParA(K20R)-CFP. Time-lapse of ChpT-YFP and ParA(K20R)-CFP dynamics within a single cell expressing *parA(K20R)* (+van), imaged at 10 min. intervals from 150 to 330 min. post-synchronization, with consecutive frames pasted side-by-side to generate the concatenated figure (left) and corresponding kymograph (right). *parA(K20R)* was induced 30 min. pre-synchronization and during imaging post-synchronization in the agarose pad by the addition of 500 μM vanillate. In the overlay, ChpT-YFP is shown in green and ParA(K20R)-CFP in red. Red arrows show the ChpT-YFP and ParA(K20R)-CFP internal cluster. Scale bar = 1 μm. (D) Bacterial two-hybrid assay. BTH101 reporter cells producing the indicated proteins or domains fused to the T18 or T25 domain of *Bordetella* adenylate cyclase were spotted on MacConkey agar plates supplemented with IPTG and maltose. Interaction between two fusion proteins results in a red color of the bacterial colony. ChpT_DHp_: ChpT Dimerization and Histidine phosphotransfer domain; CckA_HK_: CckA Histidine Kinase domain; CckA_RD_: CckA receiver domain; ChpT*: ChpT(R167E, R169E, R171E). (E) Time-lapse imaging of ChpT-sfGFP dynamics when produced (+xyl) in cells expressing *parA(K20R)* (+van), imaged at 10 min. intervals from 30 to 270 min. post-synchronization, with consecutive frames pasted side-by-side to generate the concatenated figure. *parA(K20R)* was induced 30 min. pre-synchronization and during imaging post-synchronization in the agarose pad with addition of 500 μM vanillate. *chpT-sfGFP* was induced at t=0 min. after synchronization and in the agarose pad with 0.3% xylose. Red arrow shows the ChpT-sfGFP internal cluster. Scale bar = 1 μm. (F) As in (E) but for ChpT(R167E,R169E,R171E)-sfGFP (ChpT*-sfGFP). Note that ChpT*-sfGFP still forms a single internal cluster (red arrow). Scale bar = 1 μm.

Our single-cell observations showed that (i) overexpression of wild-type *parA* led to prolonged localization of ChpT-sfGFP at the swarmer pole compared to wild-type cells (compare Fig. 4A and 4F) and (ii) in cells overexpressing *parA(K20R),* which we found prevents ChpT-sfGFP localization (Fig. 4E), ParA accumulation at the swarmer pole is reduced (Laloux and Jacobs-Wagner, 2013). Given these observations, we hypothesized that ParA may normally promote ChpT localization to the new pole in predivisional cells and provide it access to the PopZ microdomain.

Consistent with a role for ParA in recruiting ChpT, we found that cells expressing an ectopic copy of *parA(K20R)* also developed a single dynamic and non-polar internal cluster of CckA-GFP, DivL-GFP, and ChpT-sfGFP (Fig. 4D-E, S4A). This internal cluster was seen in a majority of elongated cells for each fusion protein (70% for CckA-GFP, 96% for DivL-GFP and 86% for ChpT-sfGFP with n=50 elongated cells in each case). These internal clusters arose relatively late in our time-lapse experiments, likely after additional rounds of DNA replication had initiated at the stalked pole, as evidenced by the fact that MipZ-CFP also formed internal clusters in elongated cells overexpressing *parA(K20R)* (Fig. 4C). However, whereas MipZ-CFP often accumulated in multiple internal clusters (Fig. 4C), we only ever saw the formation of a single internal cluster of CckA-GFP, DivL-GFP, and ChpT-sfGFP (Fig. 4D-E, S4A). We hypothesized that ParA(K20R) was accumulating at one of the newly formed *parS*/ParB/MipZ complexes (next to a newly replicated origin) and was recruiting CckA-GFP, DivL-GFP, and ChpT-sfGFP to this cytoplasmic position within the cell. To test this hypothesis, we expressed ParA(K20R) fused to CFP in cells also expressing YFP-MipZ or ChpT-YFP. We found that ParA(K20R)-CFP accumulated in a single internal cluster that colocalized with one of the YFP-MipZ internal clusters (Fig. S5C) and with the single ChpT-YFP internal cluster (Fig. 5C).

Previous work found that ChpT, CckA, and ParA all interact with PopZ (Schofield *et al*., 2010; Holmes *et al*., 2016). We therefore tested if PopZ was accumulating at these internal clusters and recruiting CckA and ChpT. In elongated cells expressing *parA(K20R),* we found that while PopZ-YFP accumulated strongly at the swarmer pole, very little PopZ-YFP signal was found in internal clusters (Fig. S4C), suggesting that ChpT and CckA accumulation in the internal clusters does not depend on PopZ.

### ParA not bound to DNA recruits ChpT to the swarmer pole

Our results suggest that ChpT-sfGFP gets recruited to internal *parS*/ParB/MipZ complexes only when an excess of ParA molecules accumulates at that complex (Fig. 4E-F, 5C). In wild-type stalked cells undergoing chromosome translocation, dimers of ParA-ATP coat the nucleoid and the newly formed *parS*/ParB/MipZ complex moves toward the swarmer pole through ParB-mediated hydrolysis of ParA-ATP into ParA-ADP (Ptacin *et al*., 2010; Schofield *et al*., 2010; Shebelut *et al*., 2010; Lim *et al*., 2014). ParA-ADP molecules released from the DNA are thought to relocate to the swarmer pole where they bind PopZ (Ptacin *et al*., 2014) (Fig. 3A). We hypothesized that ChpT is normally targeted to the swarmer pole or to internal clusters by binding

ParA molecules not bound to DNA. Consistent with this hypothesis, we did not observe ChpT moving across cells with the translocating chromosome in wild-type cells when ParA is mostly nucleoid-associated (Fig. 4A). To further test this hypothesis and to examine whether ParA and ChpT interact, we used a bacterial two-hybrid system in which two proteins of interest are fused to the T18 and T25 portions of adenylate cyclase. If the two proteins interact, they reconstitute adenylate cyclase activity, leading to production of a red pigment. We did not detect any interaction between ChpT and wild-type ParA (Fig. 5D). However, when testing ChpT and ParA(R195E), a DNA-binding deficient mutant of ParA (Ptacin *et al*., 2010; Schofield *et al*., 2010; Lim *et al*., 2014), we now observed colonies with intense red staining, indicative of an interaction (Fig. 5D), and supporting a model in which ChpT can bind ParA that is not bound to DNA.

ParA can dimerize and cycle through both ATP and ADP bound states. To test how these features of ParA impact interaction with ChpT, we tested different point mutants of ParA using our two-hybrid assay: ParA(G16V), a dimerization-deficient mutant that can presumably bind ATP or ADP but not DNA; ParA(K20R), an ATP-binding deficient mutant described above; and ParA(D44A), an ATP-locked mutant (Ptacin *et al*., 2010). For G16V, we still observed some red staining, suggesting that this variant still binds ChpT (Fig. S5D), likely because disrupting dimerization prevents or diminishes DNA binding by ParA. For K20R and D44A, these mutants of ParA did not interact with ChpT (Fig. S5D). However, combining each with the R195E mutation that disrupts DNA-binding produced red colonies indicative of an interaction (Fig. S5D). We conclude that the DNA-binding status of ParA, and not its ATP/ADP state, is most critical for an interaction with ChpT.

We also detected an interaction between the histidine kinase domain of CckA (CckA_HK_) and ParA(R195E), but not with wild-type ParA (Fig. 5D). Further, we found that ChpT and CckA_HK_ can each interact to some extent with ParB in our bacterial two-hybrid assay (Fig. 5D), but did not observe any interaction with either MipZ or the DNA-binding deficient mutant MipZ(R194A) (Corrales-Guerrero *et al*., 2020) (Fig. 5D). Finally, we found that the C-terminal domain of ChpT, which weakly resembles the ATP-binding domain of a histidine kinase (Fioravanti *et al*., 2012; Blair *et al*., 2013), was required for its interaction with ParA and ParB (Fig. 5D). The N-terminal domain of ChpT, which resembles the DHp domain found in histidine kinases, is critical for mediating phosphotransfer from CckA and to CtrA. The function of ChpT’s C-terminal domain had been unclear, although required for CckA binding (Fig. 5D), but our results now suggest it mediates binding to ParA that is not bound to the nucleoid.

Based on our bacterial two-hybrid assay, we infer that ChpT and CckA are recruited to internal clusters by excess levels of ParA that are unbound to DNA and associated with a ParB/*parS* complex. However, while ChpT recruitment to the new pole is disrupted when chromosome segregation is incomplete, CckA polarization is not (Iniesta *et al*., 2010b), likely because it depends on PodJ (Fig. 1D-E) rather than ParA (Fig. 4B, D). To confirm that ChpT was being recruited to internal clusters independent of CckA, we constructed, informed by the available ChpT structures (Fioravanti *et al*., 2012; Blair *et al*., 2013), a ChpT variant containing the substitutions R167E, R169E, and R171E, which we call ChpT*, that cannot bind CckA but retains interaction with ParA(R195E) and ParB (Fig. 5D). We then ectopically expressed either ChpT or ChpT* fused to sfGFP in cells with a vanillate-inducible copy of *parA(K20R)*. In the absence of vanillate, ChpT-sfGFP localized to both poles (Fig. S5E), but ChpT*-sfGFP did not localize at either cell pole at any stage of the cell cycle (Fig. S5F). These results support the notion that ChpT interaction with CckA is necessary but not sufficient for ChpT to accumulate at the poles. In the presence of vanillate to induce *parA(K20R),* ChpT-sfGFP accumulated at the old, stalked pole and in an internal cluster (Fig. 5E). By contrast, ChpT*-sfGFP accumulated only in an internal cluster (Fig. 5F). These results demonstrate that ChpT recruitment to internal clusters occurs in a CckA-independent manner and likely reflects its direct interaction with the chromosome segregation machinery. Taken all together, our results support a model in which chromosome translocation normally leads to an accumulation of ParA unbound to DNA at the nascent swarmer pole, which recruits ChpT to enable phosphotransfer from CckA to CtrA and, ultimately, successful completion of the cell cycle.

## Discussion

### A substrate-product relationship that couples cell division to DNA replication

How organisms ensure the correct order of events during the cell cycle is critical to their survival. Hartwell and Weinert outlined two general mechanisms for enforcing order (Hartwell and Weinert, 1989). One involves dedicated checkpoints, which they envisioned as surveillance systems that are not involved in executing either of two events but are involved only in monitoring and promoting their relative order of execution. The second involves substrate-product relationships in which the product of one event is the substrate for the next, thereby intrinsically ensuring they occur only in succession. Here, we uncovered a substrate-product relationship that couples cell division in *Caulobacter* to the successful initiation of DNA replication (Fig. 6). The late stages of the *Caulobacter* cell cycle include flagellar and pili biogenesis, as well as cell division. These events all depend on the phosphorylation of CtrA, which is driven by CckA and ChpT. The localization of these latter two factors to the swarmer cell pole is necessary for CtrA activation and dependent on the initiation of DNA replication (Fig. 1D-G, Fig. 6) (Iniesta *et al*., 2010b). DnaA, likely in the ATP-bound form, is a transcription factor that promotes the synthesis of PodJ (Hottes *et al*., 2005), which then recruits CckA and DivL to the swarmer pole. However, even in cells ectopically producing PodJ (and, consequently, with CckA and DivL at the pole), CtrA is not activated (Fig. 1B-C, Fig. 6). We found that ChpT is not localized unless the *ori*-proximal region of a newly replicated chromosome is properly translocated across the cell to the swarmer pole (Fig. 4E, Fig. 6). Our findings suggest that ParA, which is nucleoid-associated when promoting the directional movement of the *parS*/ParB complex, eventually accumulates at the pole and is not bound to DNA, enabling it to recruit ChpT and complete the CckA-ChpT-CtrA phosphorelay (Fig. 6). In this manner, the translocating chromosome is a physical cue that ultimately triggers CtrA activation, thereby ensuring that cell division, and other CtrA-dependent processes, do not occur until after DNA replication has successfully initiated.

**Figure 6:**
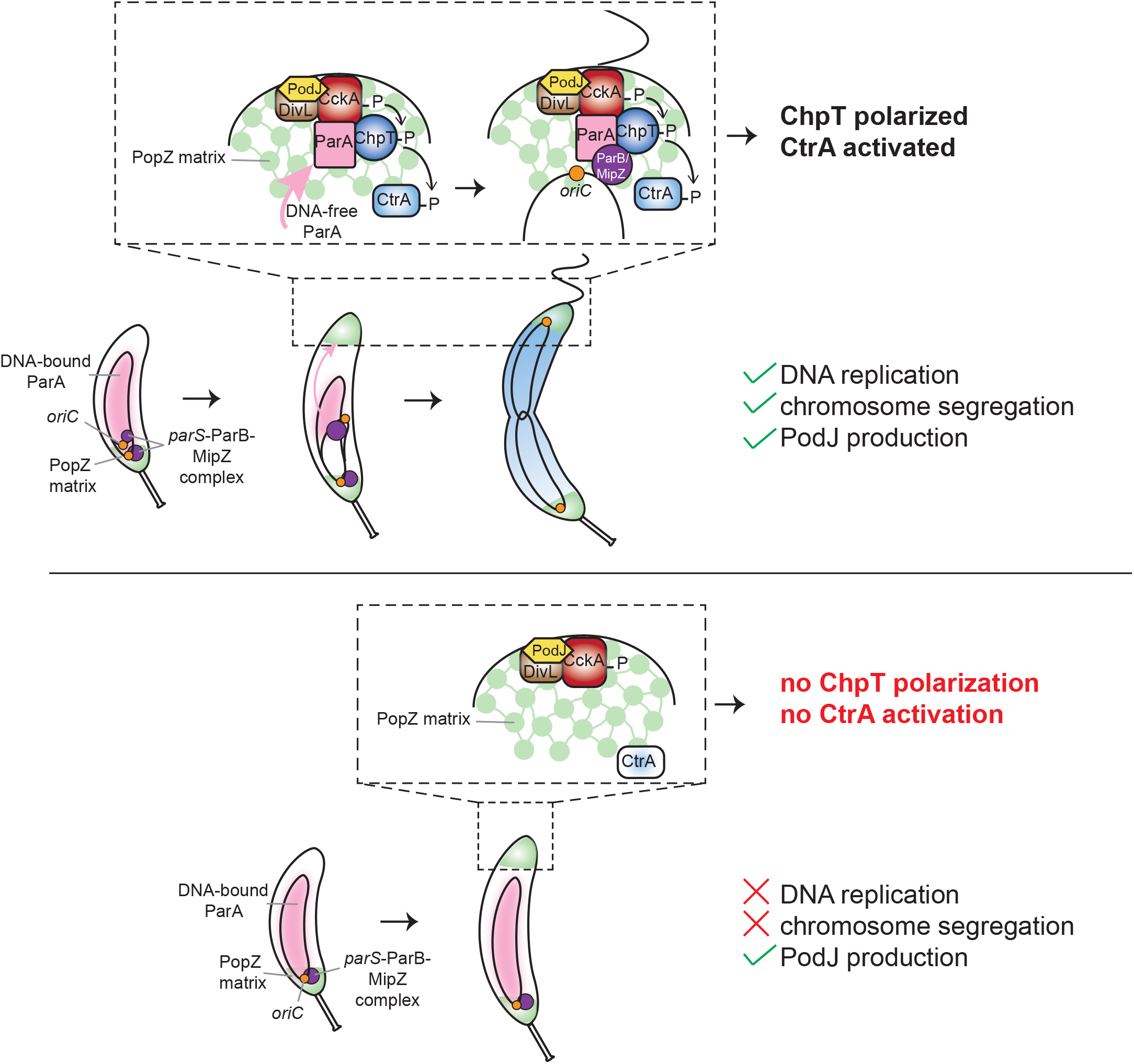
Model for coupling DNA replication initiation and proper chromosome translocation to the activation of the CckA-ChpT signaling pathway. Schematic of the polarization of CtrA regulators at the new pole. In wild-type conditions (top panel), initiation of DNA replication and proper accumulation of ParA at the new pole within the PopZ matrix enables accumulation of the ChpT intermediate component of the CckA-ChpT-CtrA phosphorelay. In cells that do not initiate DNA replication (bottom panel), production of the scaffold protein PodJ can restore the polarization of CckA and DivL, but does not allow ChpT accumulation within the PopZ matrix, leading to an incomplete pathway missing the intermediate ChpT histidine phosphotransferase for CtrA activation.

The use of chromosome position to couple cell-cycle processes also arises in many species to tie the final stages of chromosome segregation to the onset of cytokinesis. For instance, in *E. coli*, unsegregated chromosomes at mid-cell accumulate the cell division inhibitor SlmA such that cytokinesis cannot occur until after chromosomes are properly separated away from mid-cell (Bernhardt and De Boer, 2005). Noc in *B. subtilis* (Wu and Errington, 2004) and MipZ in *C. crescentus* (Thanbichler and Shapiro, 2006) play analogous roles. Notably, these systems for coupling chromosome segregation to cell division and the mechanism identified here for coupling replication to cell division feature substrate-product relationships, rather than checkpoint mechanisms involving regulatory feedback systems, as is commonly found in eukaryotes. Understanding why bacteria may rely more heavily on substrate-product relationships to order their cell cycles remains a fascinating question for the future.

### Activation of a signaling pathway by the physical translocation of a chromosome

For most two-component signaling pathways, the signals that activate them remain unknown. For cases in which a signal has been identified, most feature small molecules or low molecular weight proteins that bind to a periplasmic or transmembrane domain of a top-level histidine kinase. Although CckA and its activator DivL both reside in the inner membrane, no periplasmic or extracellular signals have been identified that regulate their activities. DivL is directly regulated by the cytoplasmic response regulator DivK (Tsokos *et al*., 2011) and CckA by cytoplasmic c-di-GMP (Lori *et al*., 2015). Our work here now indicates that a major cue for the CtrA phosphorelay involves the initially duplicated region of the chromosome, which serves to recruit ChpT via the ParA and ParB segregation complex. There is some precedent for signal sensing by intermediate components of complex phosphorelays, with Rap phosphatases in *B. subtilis* known to regulate the response regulator Spo0B, which shuttles phosphoryl groups from KinA/B/C/D/E to the histidine phosphotransferase Spo0F (Solomon *et al*., 1996; Perego, 2013). However, in that system, the Rap phosphatases respond to small Phr peptides, not a physical cue like chromosome translocation.

Notably, our work indicates that CckA alone is not sufficient to recruit ChpT to the swarmer pole, despite being sufficient to bind and phosphorylate ChpT *in vitro* (Biondi *et al*., 2006). This difference likely reflects a constraint, or barrier, imposed by the PopZ matrix that is established at the swarmer cell pole. PopZ forms a dense, interconnected matrix with possible phase-separated properties that enable the compartmentalization of the cytoplasm (Bowman *et al*., 2010; Schofield *et al*., 2010). We found that ChpT cannot access this polar microdomain unless translocation of the *parS*/ParB complex has occurred, likely because the movement of this complex across the cell is required to produce a pool of ParA at the pole that is not nucleoid bound. Although CckA is not sufficient to recruit ChpT inside the PopZ matrix, ChpT interaction with CckA is required for its accumulation at the nascent swarmer pole.

We found that in a bacterial two-hybrid system, ChpT and CckA could interact with ParA, but only if ParA was not DNA bound. Additionally, we found that in *Caulobacter* cells in which chromosome translocation was disrupted, ChpT and CckA were recruited to internal sites within the cytoplasm harboring the *parS*/ParB/ParA machinery. However, only ChpT access to the swarmer pole was dependent on a successful segregation of the chromosome; CckA accumulation at the swarmer pole was not. Importantly, ChpT co-localization with the Par machinery at internal clusters occurred even for a ChpT variant engineered to ablate binding to CckA, supporting the notion that ChpT recruitment by ParA and ParB is independent of CckA. Whether ChpT binds simultaneously to both ParA/ParB and CckA/CtrA at the swarmer pole is not yet clear. Further work is needed to explore the biochemical and biophysical details of ChpT’s multiple protein-protein interactions.

### Concluding Remarks

Our results indicate that the PopZ microdomain may serve multiple functions in *Caulobacter*. As suggested previously, it may increase the effective concentration of various signaling proteins at the pole to promote or increase their activities (Bowman *et al*., 2010; Holmes *et al*., 2016; Lasker *et al*., 2020). In addition, we propose that it also serves as a gatekeeper that selectively excludes some proteins like ChpT until others have arrived, thereby providing regulatory capabilities to cells. In a similar way, phase-separated granules and bodies in the cytoplasm of eukaryotic cells are emerging as powerful regulators and organizers of many cellular processes (Banani *et al*., 2017; Gomes and Shorter, 2019).

The mechanism identified here in which chromosome translocation serves as a physical cue to trigger a signaling pathway is likely conserved. Homologs of ChpT, PopZ, and the *parS*/ParB/ParA system are found throughout the α-proteobacteria and could work in similar ways. The Par system is found even more broadly and it has become increasingly clear that ParB and ParA can interact with a wide variety of proteins other than each other (Thanbichler and Shapiro, 2006; Gruber and Errington, 2009; Schofield *et al*., 2010; Mercy *et al*., 2019; Kawalek *et al*., 2020; Pióro and Jakimowicz, 2020), and so could participate in the regulated recruitment of proteins in other contexts and for other purposes.

In sum, our work reveals a new mechanism by which bacterial cells coordinate two cell-cycle events, ensuring that cell division does not inadvertently occur before DNA replication in *C. crescentus*. Although the timing and execution of each individual process – DNA replication initiation and cell division – has been extremely well-studied, how cells sense whether the former has occurred and relay this information to the latter has remained unclear. We found a novel connection linking the physical translocation of the origin-proximal region of a chromosome to the activation of a signaling pathway that controls cell-cycle progression. Chromosome movement and positioning is a common signal in eukaryotes; for instance, the failure to properly segregate chromosomes activates the spindle checkpoint pathway (Gorbsky, 2015). Our work suggests that bacteria may similarly use chromosome dynamics as an important cue for regulating cell-cycle progression.

## Acknowledgments

We thank S. Srikant, K. Gozzi, M. Guo, and C. Tsokos for comments on the manuscript, and C. Tsokos and J. Dyer for early discussions on the project. Instrumentation resources from the BioMicro Center in the Department of Biology at MIT are gratefully acknowledged. This work was supported by a long-term fellowship (LT000322/2017-L) from the Human Frontier Science Program to M.G. and an NIH grant to M.T.L. (R01GM082899), who is also an Investigator of the Howard Hughes Medical Institute.

## Author Contributions

M.G. performed all experiments, with assistance from A.G.S and L.K.C. M.G. and M.T.L. designed experiments, analyzed data, prepared figures, and wrote the manuscript.

## Declaration of Interests

The authors declare no competing interests.

## Methods

### Growth conditions

*Caulobacter crescentus* strains were grown in PYE (2 g/L bactopeptone, 1g/L yeast extract, 0.3 g/L MgSO_4_, 0.5 mM 0.5M CaCl_2_) or in fresh M2G+ (0.87 g/L Na_2_HPO_4_, 0.53 g/L KH_2_PO_4_, 0.5 g/L NH_4_Cl, 0.5 mM MgS0_4_, 10 µM FeS0_4_, 0.1 mM CaCl_2_, 1% PYE, 0.2% glucose) at 30°C. Expression from the P*_lac_* promoter was induced with 1 mM IPTG. Expression from the P*_xyl_* promoter was repressed with glucose (0.2%) and induced with xylose (0.3%) unless noted. Expression from the P*_cumate_* promoter was induced with 0.2 µM cumate. Expression from the P*_van_* promoter was induced with vanillate (500 µM) unless noted. To maintain plasmids, antibiotics were added at the following concentrations (liquid/plate): kanamycin (5 μg mL^-1^ / 25 μg mL^-1^), oxytetracycline (1 μg mL^-1^ / 2 μg mL^-1^), gentamycin (2.5 μg mL^-1^ / 5 μg mL^-1^), chloramphenicol (2 μg mL^-1^ / 1 μg mL^-1^).

*E. coli* strains were grown in LB (10 g/L NaCl, 10 g/L tryptone, 5 g/L yeast extract) and supplemented with antibiotics at the following concentrations unless noted (liquid/plate): kanamycin (30 μg mL^-1^ / 50 μg mL^-1^), oxytetracycline (12 μg mL^-1^ / 12 μg mL^-1^), gentamycin (15 μg mL^-1^ / 20 μg mL^-1^), chloramphenicol (20 μg mL^-1^ / 30 μg mL^-1^), carbenicillin (50 μg mL^-1^ / 100 μg mL^-1^).

### Strain construction

All *Caulobacter* strains were derivative of the wild-type isolate ML76 from the CB15N/NA1000 strain. Strains ML3364 and ML3365 were constructed using ΦCr30 phage transduction (Ely, 1991) to move pMT585-*gcrA*-3xFLAG integrated at the *xyl* locus from ML2297 into ML2000 and selecting on kanamycin + glucose + IPTG. This intermediate strain was then electroporated with pRVMCS-5-*ctrA* or pRVMCS-5-*ctrA*Δ3Ω respectively selecting on tetracycline + kanamycin + glucose + IPTG.

To construct ML3361 and ML3362, ML2000 was first transduced using ΦCr30 phage transduction (Ely, 1991) with ML1681 or ML1756 respectively and selected on gentamycin + IPTG plates. These intermediate strains were then electroporated with pQF-*podJ* and selected on oxytetracycline + gentamycin + IPTG. Strain ML3378 was constructed by electroporating *p*BXMCS-2-*dnaA(R357A)* into the same intermediate strain as above from ML2000’s transduction with ML1681 and selecting on kanamycin + IPTG + glucose plates. Strain ML3379 was constructed by electroporating *p*RVMCS-5-*ctrA* into ML3378 and selecting on oxytetracycline + kanamycin + IPTG + glucose plates.

ML1071 is a lab collection strain. Although the details of this construction are unknown, the linker between ChpT and the C-terminal YFP was sequenced and contains the following sequence: CGGGGCTCGGGCGCCACCATG.

ML2000 was transformed with pQF-*podJ* by electroporation followed by selection on oxytetracycline + IPTG. This intermediate strain was used to construct: ML3363 using ΦCr30 phage transduction (Ely, 1991) to move ChpT-YFP from ML1071; ML3359 and ML3360 by electroporating *p*JS14 P*_xylX_::ctrA* or *p*JS14 P*_xylX_::ctrA*Δ3Ω, respectively and selecting on oxytetracycline + chloramphenicol + glucose + IPTG plates; ML3367 by electroporating *p*BVMCS-4-*PA5295* and selecting on oxytetracycline + gentamycin + IPTG plates.

To construct ML3370, ML3372 and ML3379, strains ML2395, ML1756 and ML1900 were electroporated with *p*RVMCS-6-*parA(K20R).* To construct ML3371, ML3382 and ML3383, ML76 was electroporated with *p*RVMCS-6-*parA(K20R)* selecting on chloramphenicol plates, and this intermediate strain was then transduced with CckA-eGFP from ML1681 for ML3371, or electroporated with *p*XGFPC-2-*chpT-sfgfp* or *p*XGFPC-2-*chpT(R167E)(R169E)(R171E)-sfgfp* and selected on chloramphenicol + kanamycin + glucose plates, to construct ML3382 and ML3383 respectively.

Strain ML3373 was constructed by electroporating *p*YFPC-2-*chpT-sfgfp* into ML76 and selecting on kanamycin plates. Strains ML3374 and ML3375 were constructed by electroporating *p*RVMCS-6-*parA(K20R)* or *p*RVMCS-6-*parA*, respectively, into ML3373.

Strain ML3376 was constructed by electroporating *p*RVMCS-6-*parA* into ML2395 and selecting on chloramphenicol + gentamycin plates.

Strains ML3377 and ML3378 were constructed by electroporating *p*RVMCS-6-*parA(K20R)-cfp* into ML3415 and ML1071 respectively.

Strains ML3380 and ML3381 were constructed by electroporating *p*BXMCS-4-*popZ* and *p*BXMCS-2-*popZ* into ML3374 and ML3371 respectively and selecting on chloramphenicol + gentamycin + kanamycin plates.

To construct ML3384, TLS1629 was transduced with ML2397 (MipZ-CFP, kan^R^) and selected on kanamycin + chloramphenicol plates.

### Time courses and synchronization

Cells were grown overnight in PYE with appropriate antibiotics and inducers (1 mM IPTG for P*_lac_*-*dnaA* strains) or repressors (0.2% glucose for xylose inducible constructions). Cells were diluted in M2G+ supplemented with appropriate antibiotics and inducers when necessary and grown to OD_600_ ∼ 0.1 to 0.4. For synchronization, cells were first centrifuged for 10 min. at 10,000 *g*. G1/swarmer cells were isolated using Percoll (GE Healthcare) density gradient centrifugation. Briefly, pellets were resuspended in equal amounts of M2 buffer (0.87 g/liter Na_2_HPO_4_, 0.53 g/liter KH_2_PO_4_, 0.5 g/liter NH_4_Cl) and Percoll and centrifuged at 10,000 *g* for 20 min. The upper ring was aspirated, and the lower ring, corresponding to swarmer cells, was transferred into a new 15-ml Falcon tube. Swarmer cells were washed in 13 ml of M2 buffer and centrifuged for 5 min. at 10,000 *g*. The pellet was resuspended in 2 ml M2 buffer and centrifuged for 1 min. at 21,000 *g*. Cells were released into M2G+ supplemented with appropriate antibiotics (and adjusted to OD_600_ ∼ 0.05 to 0.2 when necessary) and the culture was split into different flasks with different inducers. At the indicated time points, samples were harvested from each flask for flow cytometry (0.15 ml), immunoblotting (1 ml), and RNA extraction followed by qRT-PCR (2 ml). For flow cytometry, samples were stored in 30% ethanol at 4°C. For immunoblotting and RNA extraction, cells were centrifuged for 1 min. at 15,000 rpm, aspirated, and frozen in liquid nitrogen.

For the DnaA depletion strains, prior to synchronization, at OD_600_ ∼ 0.1 to 0.25, cells were centrifuged at 10,000 *g* for 10 min, washed 2 times with M2G+, released into M2G+ with appropriate antibiotics without IPTG for 1.5 hours. Cells were then synchronized as described above.

For cells induced presynchronization, the culture was split into two flasks and one was induced with 500 µM vanillate for the indicated time. OD_600_ before induction were adjusted to be less than 0.4 at the end of the induction time. Cells were then synchronized as described above.

### Flow cytometry

Flow cytometry was performed as described previously (Guzzo *et al*., 2020). A fraction of fixed cells from the time course sampling (corresponding to an OD_600_ of ∼ 0.005) were centrifuged at 6,000 rpm for 4 min. Pelleted cells were resuspended in 1 ml of Na_2_CO_3_ buffer containing 3 µg/ml RNase A (Qiagen) and incubated at 50°C for at least 4 h. Cells were supplemented with 0.5 µl/ml SYTOX Green nucleic acid stain (Invitrogen) in Na_2_CO3 buffer and analyzed on a MACSQuant VYB flow cytometer.

### Reverse transcription coupled to quantitative PCR

Reverse transcription coupled to quantitative PCR was performed as described previously (Guzzo *et al*., 2020). RNA was extracted using hot TRIzol lysis and the Direct-zol RNA miniprep kit (Zymo). 2.5-µl of RNA at 50-100 ng/µl was mixed with 0.5 µl of 100-ng/µl random hexamer primers (Invitrogen), 0.5 µl of 10 mM deoxynucleoside triphosphates (dNTPs) and 3 µl of diethylpyrocarbonate (DEPC) water; incubated at 65°C for 5 min; and then placed on ice for 1 min. Two microliters of first-strand synthesis buffer, 0.5 µl of 100 mM dithiothreitol (DTT), 0.5 µl of SUPERase-In (ThermoFisher), and 0.5 µl of Superscript III (ThermoFisher) were added to each tube, and the following thermocycler program was used: 10 min. at 25°C, 1 h at 50°C, and 15 min. at 70°C. One microliter of RNase H (NEB) was added to each tube, and each reaction mixture was incubated at 37°C for 20 min.

cDNA solutions were diluted 10 times in nuclease-free water for quantitative PCR (qPCR). One microliter of diluted cDNA or serially diluted genomic DNA (gDNA) used as a standard curve was mixed with an appropriate pair of primers, i.e., either rpoA_qPCR_1 and rpoA_qPCR_2 as a control, ctrA_qPCR_1 and ctrA_qPCR_5, divK_qPCR_1 and divK_qPCR_2, pdeA_qPCR_3 and pdeA_qPCR_5 or podJ_qPCR_1 and podJ_qPCR_2, and 2X qPCR Master Mix. All experimental samples were loaded as duplicates and with standard curves on a 384-well plate for qPCR. qPCR was conducted in a LightCycler 480 system (Roche) using the following thermocycler program: 95°C for 10 min, 95°C for 15 s, 60°C for 30 s, and 72°C for 30 s with 40 cycles of steps 2 to 4.

### Immunoblotting

Immunoblotting was performed as described previously (Guzzo *et al*., 2020). Frozen pellets from the time course sampling were normalized by OD_600_ for resuspension in 1x blue loading buffer (NEB) supplemented with 1x reducing agent (DTT), boiled at 95°C for 10 min, and loaded on 12% gels (Bio-Rad) for electrophoresis. Proteins were transferred from the gel into polyvinylidene difluoride (PVDF) membranes and immunoblotted. Antibodies were used at the concentrations shown in parentheses: anti-RpoA (1:5,000, BioLegend), anti-CtrA (1:5,000), anti-SciP (1:2000) and anti-GcrA (1:5,000). Horseradish peroxidase (HRP)-conjugated secondary antibodies (ThermoFisher) were used at the concentrations shown in parentheses: anti-mouse (1:10,000) and anti-rabbit (1:5,000). The membranes were developed with SuperSignal West Femto maximum-sensitivity substrate (ThermoFisher) and visualized with a FluorChem R Imager (ProteinSimple).

### Microscopy

Fixed cells: Cells were grown overnight in PYE with appropriate antibiotics and 1 mM IPTG. Cells were diluted in M2G+ supplemented with appropriate antibiotics and 1 mM IPTG and grown to OD_600_ ∼ 0.1 to 0.25. DnaA was depleted and cells were synchronized as described above (see Time courses and synchronization section). Cells were released into M2G+ supplemented with appropriate antibiotics (and adjusted to OD_600_ ∼ 0.05 to 0.2 when necessary) and the culture was split into different flasks with different inducers. At t=90 min, 1 mL of culture for each condition was harvested and 25 µl of 16% Paraformaldehyde Aqueous Solution (Fisher Scientific) was added to the cells and gently mixed by inversion. After 1 min. of fixation at room temperature, cells were centrifuged for 1 min. at 10,000 *g*. Pelleted cells were washed once in 1 x PBS and then resuspended in 1 x PBS. One microliter of cells was spotted onto PBS-1.5% agarose pads and imaged. Phase-contrast and epifluorescence images were taken on a Zeiss Observer Z1 microscope using a 100x/1.4 oil immersion objective and an LED-based Colibri illumination system using MetaMorph software (Universal Imaging, PA).

Live imaging: Cells were grown overnight in PYE with appropriate antibiotics and repressor when appropriate (0.2% glucose for xylose inducible constructions). Cells were diluted in M2G+ supplemented with appropriate antibiotics and grown to OD_600_ ∼ 0.1 to 0.3. For cells induced presynchronization, the culture was split into two flasks and one was induced with 500 µM vanillate for 30 minutes. Cells were then synchronized as described above. After synchronization, cells were released in M2G+ with or without inducer (note that xylose when indicated is added at t=0 and not prior to synchronization). Cells were grown in flask for 10 min. and then one microliter of cells was spotted onto M2G+-1.5% agarose pads containing or not appropriate inducers in a 35 mm Glass bottom dish. Phase-contrast and epifluorescence images were taken on a Zeiss Observer Z1 microscope using a 100x/1.4 oil immersion objective and an LED-based Colibri illumination system using MetaMorph software (Universal Imaging, PA). Images were taken every 10 minutes for up to 250 minutes starting 20 minutes or 30 minutes after synchronization in a 30°C incubation chamber.

### Bacterial two-hybrid

Protein interactions were assayed using the bacterial adenylate cyclase two-hybrid system (Karimova *et al*., 1998). Genes of interest were fused to the 3’ or 5’ end of the T18 or T25 fragments of *Bordetella* adenylate cyclase using the pUT18C, pUT18, pKT25, or pKNT25 vectors. Different combinations of T18-/T25-fusion plasmids were co-transformed into the BTH101 *E. coli* strain. Co-transformants were grown in LB supplemented with kanamycin and carbenicillin overnight. Saturated cultures were spotted onto MacConkey agar (40 g/L) plates supplemented with maltose (1%), IPTG (1 mM), kanamycin (25 μg mL^-1^) and carbenicillin (50 μg mL^-1^). Plates were incubated at 30°C and pictures taken after two days.

### qPCR analysis of origin-terminus ratio

Cells were grown overnight in PYE with appropriate antibiotics, 1 mM IPTG and 0.2% glucose. Cells were diluted in M2G+ supplemented with appropriate antibiotics and 1 mM IPTG and grown to OD_600_ ∼ 0.4. To deplete DnaA, cells were centrifuged at 10,000 *g* for 10 min, washed 3 times in M2 and diluted into M2G+ with appropriate antibiotics without IPTG for 2 hours. After the 2-hour depletion, the cultures (OD_600_ ∼ 0.4) were synchronized as described above. After synchronization, cells were released into M2G+ and split in different flasks with the indicated inducers. Samples were harvested at the indicated timepoints for genomic DNA (gDNA) extraction. gDNA was extracted by resuspending pellets in 600 μL Cell Lysis Solution (QIAGEN) and incubating at 80°C for 5 min. to lyse cells. RNAs were removed by treatment with 50 μg RNase A (QIAGEN) at 37°C for 30 min. 200 μL Protein Precipitation Solution (QIAGEN) was added, the sample vortexed, and left on ice for 30 min. to precipitate proteins. After spinning at 14,000 rpm for 10 min, the supernatant was transferred to a tube containing 600 μL isopropanol and mixed by inversion. DNA was harvested by spinning at 14,000 rpm for 1 min. followed by a wash with 600 μL of 70% ethanol. The DNA pellet was resuspended in 50-100 μL H_2_O. For qPCR, DNAs were diluted 1:500 and mixed with either Cori_2 (chromosomal position: 0 Mb), CCNA_03518 (chr. pos.: 3.676 Mb), CCNA_00623 (chr. pos.: 0.669 Mb), CCNA_02846 (chr. pos.: 2.999 Mb), CCNA_02567 (chr. pos.: 2.717 Mb), CCNA_02189 (2.340 Mb) or CCNA_01869 (chr. pos.: 2.008 Mb) forward/reverse primer mix and 2X qPCR Master Mix. All experimental samples and standard curves were loaded onto a 384-well plate in duplicate for qPCR. qPCR was conducted in a LightCycler 480 system (Roche) using the following thermocycler program: 95°C for 10 min, 95°C for 15 s, 60°C for 30 s, and 72°C for 30 s with 40 cycles of steps 2 to 4.

### Plasmid construction

#### Integration plasmids

*pYFPC-2-chpTsfGFP:* the pYFPC-2 plasmid (Thanbichler *et al*., 2007) was amplified using primers pYFPC2_chpT_up_R and sfGFP_dwn_pYFPC_2_F and the *chpT* gene was amplified using primers pYFPC2_chpT_up_F and chpT_dwn_sfGFP_up_R. The *chpT* PCR product, a *sfGFP* gblock codon optimized for *Caulobacter* from IDT, and the PCR fragment of the plasmid were assembled together using the Hifi DNA assembly mix (NEB). The construction was verified by sequencing.

*pXGFPC-2-chpT-sfGFP*: The *chpT-sfGFP* insert was amplified from pYFPC-2-*chpT-sfgfp* using primers pBX_chpT_up_F and sfGFP_pXGFPC_down_R2. The pXGFPC-2 plasmid (Thanbichler *et al*., 2007) was amplified using primers sfGFP_pXGFPC_down_F2 and pBX_chpT_up_R. The two PCR products were assembled together using the Hifi DNA assembly mix (NEB). The construction was verified by sequencing.

*pXGFPC-2-chpT(R167E)(R169E)(R171E)-sfGFP*: same cloning protocol as pXGFPC-2-*chpT-sfGFP* but using *p*KNT25-ChpT(R167E)(R169E)(R171E)-T25 as a template. The construction was verified by sequencing.

#### Replicative plasmids

*P_cumate_-podJ:* The *podJ* insert was amplified from a P*_van_*-*podJ* containing plasmid using primers podJ_4pQF_1..24 and podJ_4pQF_last24 and. The pQF plasmid (Kaczmarczyk *et al*., 2013) was amplified using primers pQF_CPEC_up and pQF_CPEC_noFLAG_down. The two PCR products were assembled together using circular polymerase extension cloning (CPEC). The construction was verified by sequencing.

*pRVMCS-5-ctrA :* This plasmid was cloned by first amplifying *ctrA* from genomic DNA using primers ctrA-NdeI-F and ctrA-extrastop-NheI-R. The PCR product was digested with NdeI and NheI-HF and ligated into similarly digested pRVMCS-5 plasmid (Thanbichler *et al*., 2007) using T4 DNA ligase. The construction was verified by sequencing.

*pRVMCS-5-ctrAΔ3Ω:* This plasmid was cloned using the same protocol as for *pRVMCS-5-ctrA* but by amplifying *ctrA* from ML46 using primers ctrA-NdeI-F and ctrAD3W-NheI-R. The construction was verified by sequencing.

*pBXMCS-2-dnaA(R357A):* This plasmid was cloned by first using around-the-horn PCR to introduce the R357A mutation into *dnaA* (from a P*_van_*-*dnaA* containing plasmid) using primers dnaA 1068..1048 and dnaA 1069..1095 R357A. The *dnaA(R357A)* allele was then amplified using primers DnaA_rev_3xTAA_EcoRI and NdeI_DnaA_1..23 and cloned into pBXMCS-2 (Thanbichler *et al*., 2007) using EcoRI/NdeI. The construction was verified by sequencing.

*pRVMCS-6-parA(K20R):* genomic DNA from ML2414 (Badrinarayanan *et al*., 2015) was used as a template to amplify *parA(K20R)* using primers pXYFP_ParA_NdeI_up_F and ParA_TAATAA_SacI_R. The PCR product was digested with NdeI and SacI restriction enzymes and ligated into similarly digested *p*RVMCS-6 plasmid (Thanbichler *et al*., 2007) using T4 DNA ligase. The construction was verified by sequencing.

*pRVMCS-6-parA:* the *parA* gene was amplified from *p*KT25-*parA* using primers pRVMCS6_parA_up_F and pRVMCS-6_parA_dwn_R and the pRVMCS-6-*parA(K20R)* plasmid template was amplified in two separate fragments using primers pRVMCS6_parA_dwn_F and pRVMCS6_upcat_R, and pRVMCS6_parA_up_R and pRMCS_mid_F. The *parA* fragment and the two plasmid fragments were assembled using the Hifi DNA assembly mix (NEB). The construction was verified by sequencing.

*pRVMCS-6-parA(K20R)-CFP:* the *parA(K20R)* fragment was amplified from pRVMCS-6-*parA(K20R)* using primers pRVMCS-6-parA_up_F and parA_CFP_R. The *cfp* fragment was amplified from pXCFPC-6 (Thanbichler *et al*., 2007) using primers parA_CFP_F and CFP_pRVMCS-6_dwn_R. The pRVMCS-6-*parA(K20R)* plasmid template was amplified in two separate fragments using primers and CFP_pRVMCS-6_dwn_F and pRVMCS6_upcat_R, and pRVMCS6_parA_up_R and pRMCS_mid_F. The *parA(K20R)* fragment, the *cfp* insert and the two plasmid fragments were assembled using the Hifi DNA assembly mix (NEB). The construction was verified by sequencing.

pBXMCS-2-*popZ* and pBXMCS-4-*popZ*: The *popZ* insert was amplified from genomic DNA using pBX_popZ_up_F and pB_popZ_down_R and the pBXMCS-2 or pBXMCS-4 (Thanbichler *et al*., 2007) plasmids were amplified using primers pB_popZ_down_F and pBX_popZ_up_R. The *popZ* insert and the plasmid PCR product were assembled together using the Hifi DNA assembly mix (NEB). The construction was verified by sequencing.

#### Bacterial two-hybrid plasmids

All bacterial two-hybrid plasmids were constructed using the Gibson assembly mix (NEB) or the Hifi DNA assembly mix (NEB). All constructions (gene of interest and T18- or T25-fragments) were verified by sequencing.

*p*KNT25-ChpT : The *chpT* fragment was amplified using primers pKNT25_chpT_up_F and pKNT25_chpT_down_R. The pKNT25 plasmid was amplified using primers pKNT25_chpT_up_R and pKNT25_chpT_down_F.

*p*KNT25-ChpT_DHP_ : The *chpT_DHP_* fragment was amplified using primers pKNT25_chpT_up_F and pKNT25_chpTDHP_down_R. The pKNT25 plasmid was amplified using primers pKNT25_chpT_up_R and pKNT25_chpTDHP_down_F.

*p*KT25-CckA_HK_: The *cckA_HK_* fragment was amplified using primers pKT25_cckA_HK_up_F and pKT25_cckA_HK_down_R. The pKT25 plasmid was amplified using primers pKT25_up_R and pKT25_down_F.

*p*KNT25-ChpT(R167E)(R169E)(R171E)-T25 : Using *p*KNT25-ChpT as a template, the plasmid was amplified using primers R167E169E171E_short2_F and R167E169E171E_short2_R. The two R167E169E171E ultramers (forward and reverse) were mixed together in equal molar amounts and duplexed using the following thermocycler program: 94°C for 3 minutes, cool down to 25°C over 45 minutes at a pace of 1.5°C per minute. The duplexed ultramers were diluted 100-fold before addition to the Gibson reaction with the plasmid fragment.

*p*UT18C-ParB : The *parB* fragment was amplified using primers pUT18C_parB_up_F and pUT18C_parB_down_R. The pUT18C plasmid was amplified using primers pUT18C_parB_up_R and pUT18C_parB_down_F.

*p*UT18C-ParA and *p*UT18C-ParA(K20R): The *parA* or *parA(K20R)* fragment were amplified using primers pUT18C_parA_up_F and pUT18C_parA_down_R. The pUT18C plasmid was amplified using primers pUT18C_parA_up_R and pUT18C_parA_down_F.

*p*UT18C-ParA(R195E): Using *p*UT18C-ParA as a template, the plasmid was amplified using primers ParAR195E_F and ParAR195E_R. The two ParA(R195E) ultramers (forward and reverse) were duplexed and used as described for *p*KNT25-ChpT(R167E)(R169E)(R171E)-T25.

*p*UT18C-ParA(K20R)(R195E): Using *p*UT18C-ParA(K20R) as a template, the plasmid was amplified using primers ParAR195E_F and ParAR195E_R. The two ParA(R195E) ultramers (forward and reverse) were duplexed and used as described for *p*KNT25-ChpT(R167E)(R169E)(R171E)-T25.

*p*UT18C-ParA(D44A): Using *p*UT18C-ParA as a template, the plasmid was amplified using primers ParAD44A_F and ParAD44A_R. The two ParA(D44A) ultramers (forward and reverse) were duplexed and used as described for *p*KNT25-ChpT(R167E)(R169E)(R171E)-T25.

*p*UT18C-ParA(D44A)(R195E): Using *p*UT18C-ParA(R195E) as a template, the plasmid was amplified using primers ParAD44A_F and ParAD44A_R. The two ParA(D44A) ultramers (forward and reverse) were duplexed and used as described for *p*KNT25-ChpT(R167E)(R169E)(R171E)-T25.

*p*UT18C-ParA(G16V): Using *p*UT18C-ParA as a template, the plasmid was amplified using primers ParAG16V_F and ParAG16V_R. The two ParA(G16V) ultramers (forward and reverse) were duplexed and used as described for *p*KNT25-ChpT(R167E)(R169E)(R171E)-T25.

*p*UT18-ChpT: The *chpT* fragment was amplified using primers pUT18_chpT_up_F and pUT18_chpT_down_R. The pUT18 plasmid was amplified using primers pUT18_chpT_up_R and pUT18_chpT_down_F.

*p*UT18C-CckA_RD_: The *cckA_RD_* fragment was amplified using primers pUT18C_cckA_RD_up_F and pUT18C_cckA_RD_down_R. The pUT18C plasmid was amplified using primers pUT18C_cckA_RD_up_R and pUT18C_cckA_RD_down_F.

*p*UT18C-MipZ: The *mipZ* fragment was amplified using primers pUT18C_mipZ_up_F and pUT18C_mipZ_down_R. The pUT18C plasmid was amplified using primers pUT18C_mipZ_up_R and pUT18C_mipZ_down_F.

*p*UT18C-MipZ(R194A): this plasmid was first constructed by inserting the R194A mutation into *mipZ* in the pKNT25-*mipZ* plasmid using primers mipZR194A_F and mipZR194A_R. The two MipZ(R194A) ultramers (forward and reverse) were duplexed and used as described for *p*KNT25-ChpT(R167E)(R169E)(R171E)-T25. The pUT18C-MipZ(R194A) plasmid was then constructed by amplifying the *mipZ(R194A)* fragment from pKNT25-*mipZ(R194A)* using primers pUT18C_mipZ_up_F and pUT18C_mipZ_down_R and the pUT18C plasmid was amplified using primers pUT18C_mipZ_up_R and pUT18C_mipZ_down_F.

### Quantification and statistical analysis

#### qRT-PCR

Crossing point (Cp) values were calculated from LightCycler 480 software at the second derivative maximum. Technical replicates were averaged to yield a final Cp value for each sample and relative quantities of cDNA were calculated based on a 3-fold dilution standard curve. Each time point value for each gene of interest was normalized to the *rpoA* measured value, as *rpoA* expression remains constant in exponential phase.

Dose response fold-differences at timepoint 100 minutes are the ratios between the normalized cDNA relative quantity of the induce condition and the non-induced condition. Independent biological replicates individual datapoints are shown as well as the average ratio.

#### qPCR/origin-ter ratio

Cp values were calculated from LightCycler 480 software at the second derivative maximum. Technical replicates were averaged to yield a final Cp value for each sample and standard curve point. Relative quantities of cDNA were calculated based on a 3-fold dilution standard curve. Chromosome copy numbers were normalized internally to CCNA_01869 (terminus region) as a loading control and then normalized to the origin abundance after synchrony (0 min) to evaluate fold-change from 1N.

#### Quantification of protein levels by immunoblotting

Image quantification and analysis was done with Fiji/ImageJ. Band intensities were measured for each protein. RpoA band intensities were used as a loading control for each sample, meaning that each protein band intensity in a specific lane were normalized to RpoA band intensity in that same lane.

#### Flow cytometry analysis

Flow cytometry was analyzed with FlowJo from 50,000 total SYTOX positive events for each experiment. For each experiment, 1N was determined based on the distribution of G1 cells and 2N based on the control condition.

#### Microscopy

Polar clusters were counted manually or using MicrobeJ. The swarmer pole was determined by identifying the stalk on phase contrast images. The percentage of cells with polar clusters at the new pole were calculated for a total number of cells as specified in the figures using at least two biological replicates for each condition. For the live microscopy, a cell was counted as containing a polar cluster at the swarmer pole or an internal cluster if a cluster was identified at any time during the course of the time-lapse. Kymographs were built using Fiji and the time lapse alignment were made using the “Straighten” function in Fiji. The fluorescence intensities from the old pole to the new pole were measured automatically using MicrobeJ. For quantifying the percentage of elongated cells in strains expressing *parA(K20R)* and MipZ, DivL, CckA, or ChpT fused to a fluorescent protein, we examined cells from two independent cultures.

## Supplemental Figure Legends

**Figure S1:**
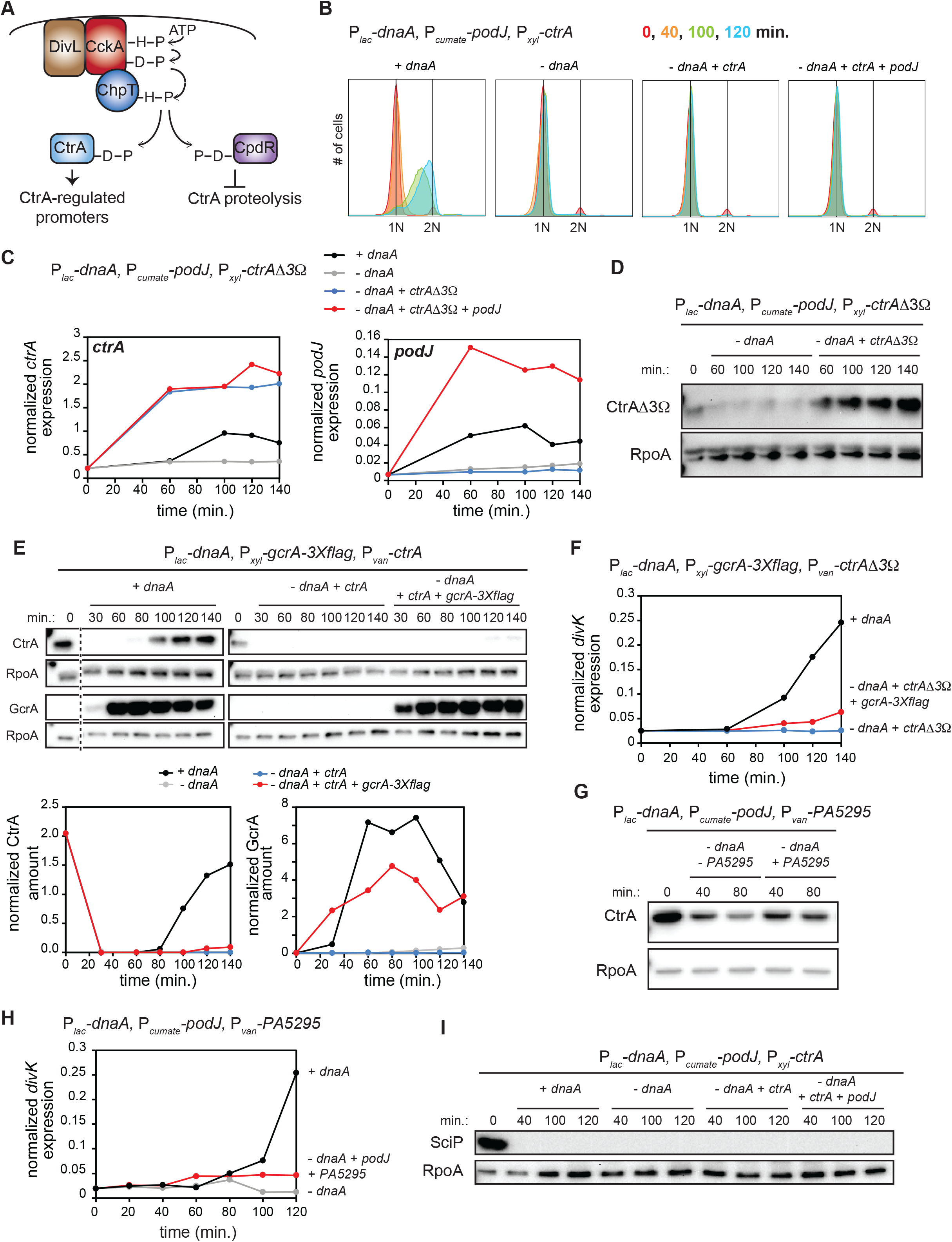
Polarization of CckA and DivL is not sufficient to activate CtrA in predivisional cells in the absence of DNA replication. (A) Schematic of the CckA-ChpT phosphorelay activity at the new pole in predivisional cells. (B) Flow cytometry profiles after SYTOX staining showing DNA content of synchronized cells expressing *dnaA* (+IPTG) or depleted of *dnaA* (-IPTG) with ectopic expression of *ctrA* (+0.075% xyl) and *podJ* (+cumate) when indicated. (C) mRNA levels of the *ctrA* and *podJ* genes measured by qRT-PCR and normalized to *rpoA* mRNA levels in cells expressing *dnaA* (+IPTG) or depleted for *dnaA* (-IPTG) with ectopic expression of the stable *ctrA* point mutant *ctrAΔ3Ω* (+0.075% xyl) and *podJ* (+cumate) when indicated. (D) CtrAΔ3Ω protein levels in cells depleted for *dnaA* (-IPTG) with ectopic expression of the stable *ctrA* point mutant *ctrAΔ3Ω* (+0.075% xyl) when indicated. (E) CtrA and GcrA protein levels at the times indicated post-synchronization in cells expressing *dnaA* (+IPTG) or depleted of *dnaA* (-IPTG) with ectopic expression of wild-type *ctrA* (+van) and *gcrA-3xflag* (+xyl) when indicated. Graphs show CtrA and GcrA protein band intensity normalized to RpoA. (F) mRNA levels of the CtrA-activated gene *divK* measured by qRT-PCR and normalized to *rpoA* mRNA levels at the times indicated post-synchronization in cells expressing *dnaA* (+IPTG) or depleted of *dnaA* (-IPTG) with ectopic expression of the proteolytically stable mutant *ctrAΔ3Ω* (+van) and with (+xyl) or without ectopic expression of *gcrA-3xflag*. (G) CtrA protein levels in cells expressing (+ 50 μM van) or not (-van) the PA5295 phosphodiesterase in cells depleted for *dnaA* (-IPTG). (H) mRNA levels of the CtrA-activated gene *divK* measured by qRT-PCR and normalized to *rpoA* mRNA levels at the times indicated post-synchronization in cells expressing *dnaA* (+IPTG) or depleted of *dnaA* (-IPTG) and the *Pseudomonas aeruginosa* phosphodiesterase PA5295 (+50 μM van) when indicated. This strain also expressed an ectopic copy of *podJ (*+cumate), when indicated. (I) SciP protein levels at the times indicated post-synchronization in cells expressing *dnaA* (+IPTG) or depleted of *dnaA* (-IPTG) with ectopic expression of *ctrA* (+xyl) and *podJ* (+cumate) when indicated.

**Figure S2:**
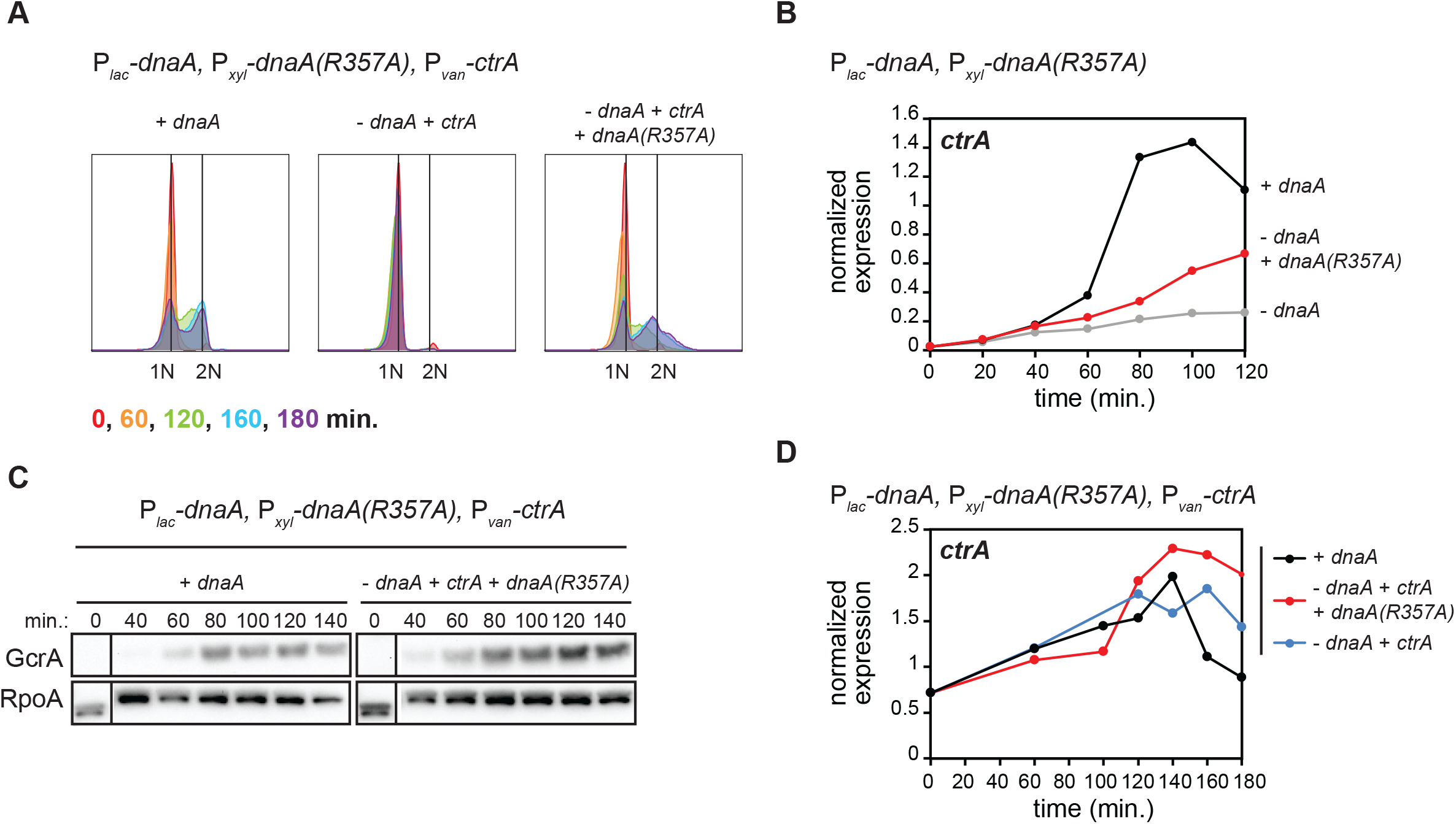
Replication of the full chromosome is not required for CtrA activation. (A) Flow cytometry profiles after SYTOX staining showing DNA content of synchronized cells expressing *dnaA* (+IPTG) or depleted of *dnaA* (-IPTG) with ectopic expression of *ctrA* (+van) and *dnaA(R357A)* (+xyl) when indicated. (B) *ctrA* mRNA levels measured by qRT-PCR and normalized to *rpoA* mRNA levels in cells expressing dnaA (+IPTG) or depleted of *dnaA* (-IPTG) with ectopic expression of *dnaA(R357A)* (+xyl) when indicated. (C) GcrA protein levels in cells expressing *dnaA* (+IPTG) or depleted of *dnaA* (-IPTG) with ectopic expression of wild-type *ctrA* (+van) and *dnaA(R357A)* (+xyl). RpoA is shown as a loading control. (D) mRNA levels of *ctrA* measured by qRT-PCR and normalized to *rpoA* mRNA levels in cells expressing dnaA (+IPTG) or depleted of *dnaA* (-IPTG) with ectopic expression of *ctrA* (+van) and *dnaA(R357A)* (+xyl) when indicated.

**Figure S3:**
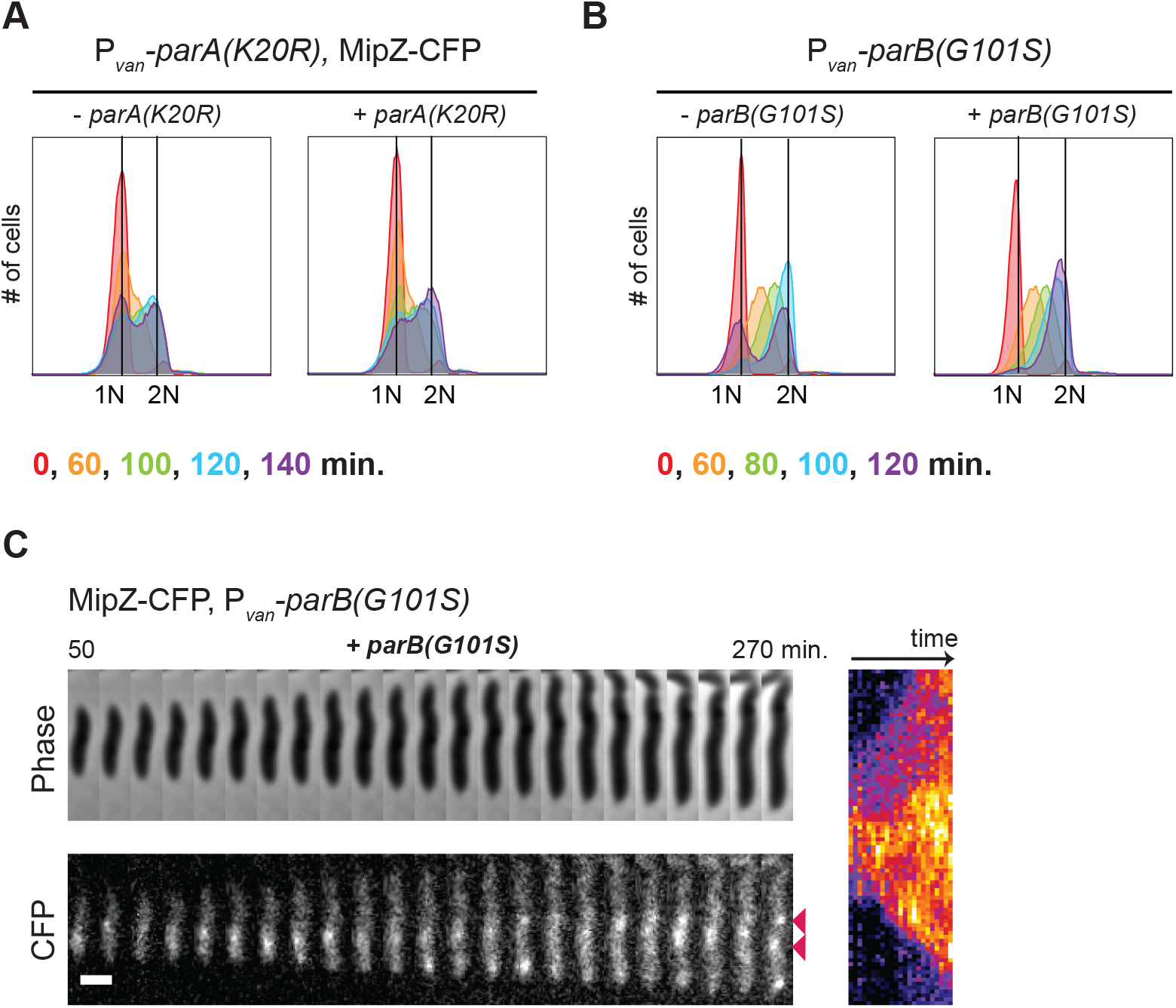
CtrA activation in predivisional cells is reduced when chromosome segregation is perturbed. (A) Flow cytometry profiles after SYTOX staining showing DNA content of synchronized cells with (+van) or without (-van) ectopic expression of *parA(K20R)*. Expression of *parA(K20R)* was induced 60 min. before synchronization and after synchronization with 500 μM vanillate. (B) Flow cytometry profiles after SYTOX staining showing DNA content of synchronized cells with (+van) or without (-van) ectopic expression of *parB(G101S)*. Expression of *parB(G101S)* was induced 60 min. before synchronization and after synchronization with 500 μM vanillate. (C) Time lapse of MipZ-CFP dynamics within a single cell expressing the *parB(G101S)* mutant, imaged at 10 min. intervals from 50 to 270 min. post-synchronization, with consecutive frames pasted side-by-side to generate the concatenated figure (left) and corresponding kymograph (right). Expression of *parB(G101S)* was induced 30 min. before synchronization and during imaging post-synchronization in the agarose pad with addition of 500 μM vanillate. The red arrows indicate MipZ-CFP internal clusters next to the newly replicated chromosomal origins. Scale bar = 1 μm.

**Figure S4:**
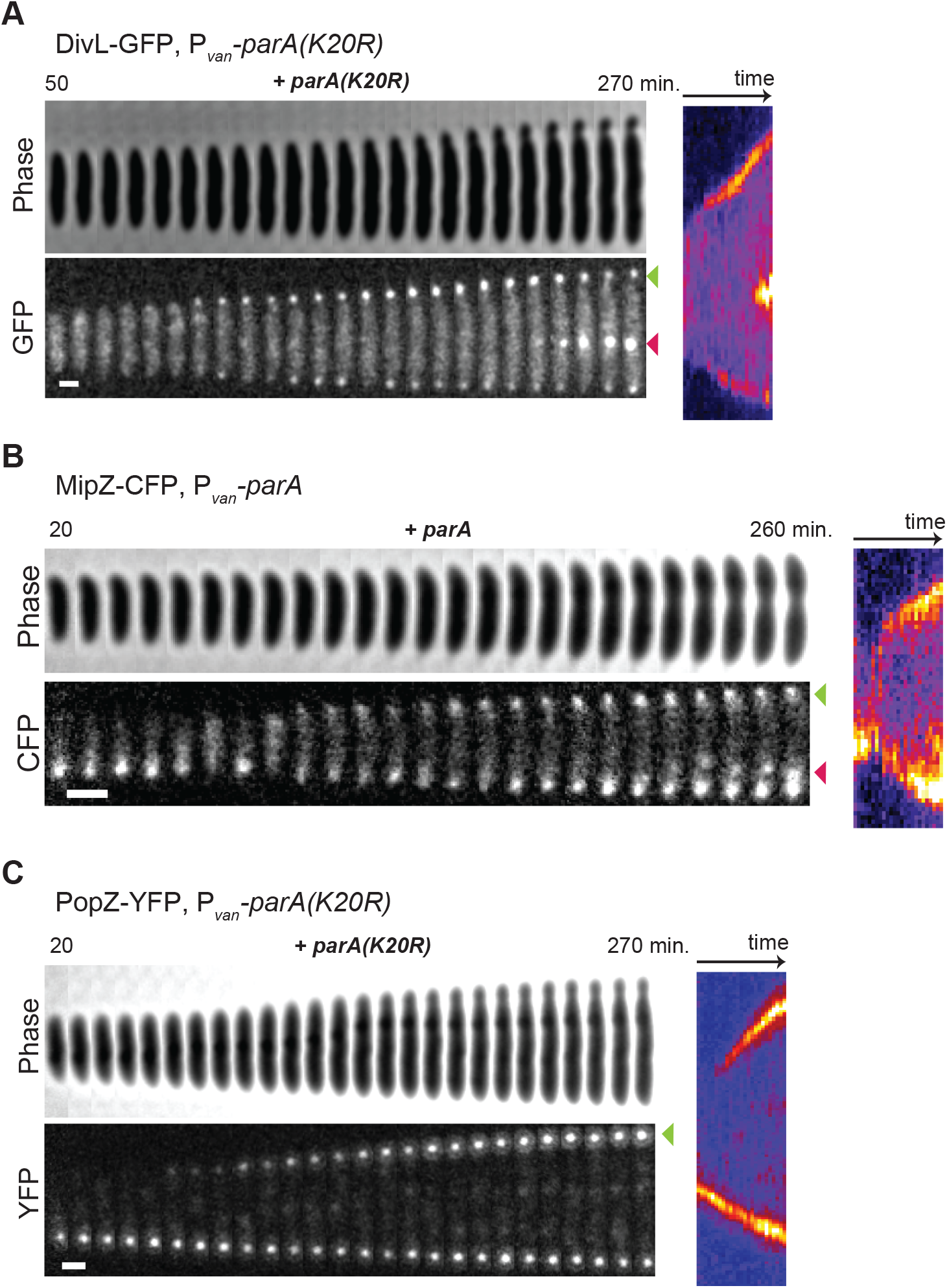
ChpT access to the new swarmer pole is affected when chromosome segregation is disrupted. (A) Time lapse of DivL-GFP dynamics within a single cell expressing the dominant negative *parA(K20R)* mutant, imaged at 10 min. intervals from 50 to 270 min. post-synchronization, with consecutive frames pasted side-by-side to generate the concatenated figure (left) and corresponding kymograph (right). Expression of *parA(K20R)* was induced 30 min. before synchronization and during imaging post-synchronization in the agarose pad with addition of 500 μM vanillate. The green arrow shows DivL-GFP localization at the new swarmer pole upon expression of *parA(K20R).* The red arrow shows DivL-GFP accumulation in a single internal cluster. Scale bar = 1 μm. (B) Timelapse of MipZ-CFP dynamics within a single cell expressing ectopic wild-type *parA*, imaged at 10 min. intervals from 20 to 260 min. post-synchronization, with consecutive timeframes pasted side-by-side to generate the concatenated figure (left) and corresponding kymograph (right). Expression of *parA* was induced 30 min. before synchronization and during imaging post-synchronization in the agarose pad with addition of 500 μM vanillate. Green arrow shows MipZ-CFP localization at the new swarmer pole upon expression of *parA.* Red arrow shows MipZ-CFP internal cluster. Scale bar = 1 μm. (C) As in (A) but for PopZ-YFP dynamics. The green arrow shows PopZ-YFP localization at the new swarmer pole upon expression of *parA(K20R).* Scale bar = 1 μm.

**Figure S5:**
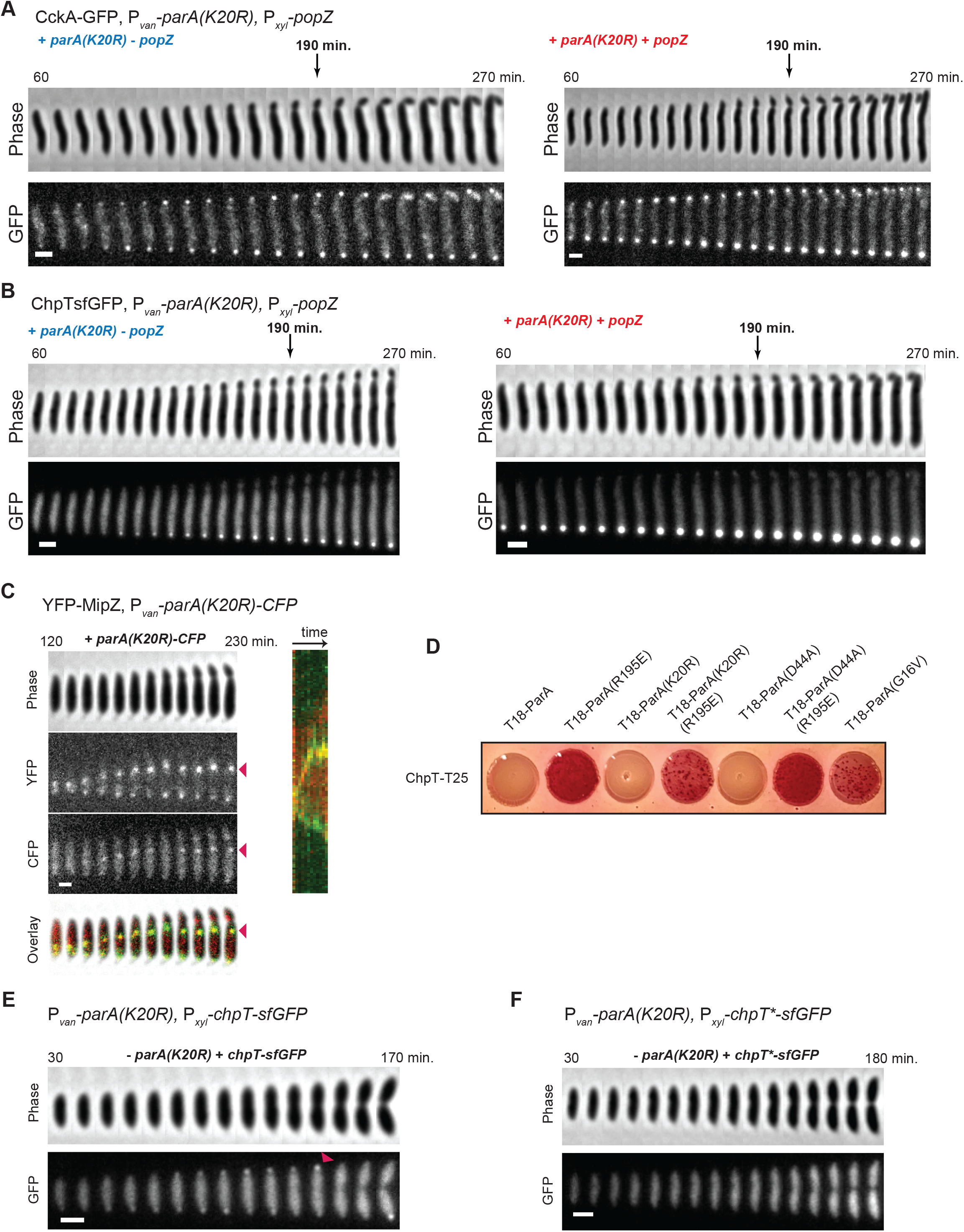
ParA not bound to DNA recruits ChpT to the swarmer pole. (A-B) Full time lapses of CckA-GFP (A) and ChpT-sfGFP (B) dynamics in cells expressing *parA(K20R)* (+van) with (+xyl) or without (-xyl) induction of *popZ* expression, corresponding to Fig. 5A and B respectively. Scale bar = 1 μm. (C) YFP-MipZ internal clusters colocalize with ectopically produced ParA(K20R)-CFP. Time lapse imaging of YFP-MipZ and ParA(K20R)-CFP dynamics within a single cell expressing *parA(K20R)* (+van), imaged at 10 min. intervals from 150 to 330 min. post-synchronization, with consecutive frames pasted side-by-side to generate the concatenated figure (left) and corresponding kymograph (right). Expression of *parA(K20R)* was induced 30 min. before synchronization and during imaging post-synchronization in the agarose pad with addition of 500 μM vanillate. In the overlay, YFP-MipZ is shown in green and ParA(K20R)-CFP in red. Red arrows show the YFP-MipZ and ParA(K20R)-CFP internal cluster. Scale bar = 1 μm. (D) Bacterial two-hybrid assay testing the interaction of ChpT fused to the T25 *Bordetella* adenylate cyclase domain with ParA and ParA point mutants fused to the *Bordetella* T18 adenylate cyclase domain. When spotted on MacConkey agar plates supplemented with IPTG and maltose, interactions between the two fusion proteins result in a red color shift of the bacterial colony. (E-F) Time lapse of the dynamics of ectopic ChpT-sfGFP and the ChpT(R167E,R169E,R171E)-sfGFP (ChpT*-sfGFP) mutant (+xyl) in cells that are not expressing *parA(K20R).* Compare to Fig. 5E and F, respectively. Note that the ChpT*-sfGFP point mutant fails to accumulate at the new pole prior to cell division, compared to ectopically produced wild-type ChpT-sfGFP (red arrow). Scale bar = 1 μm.

## Notes

### Competing Interest Statement

The authors have declared no competing interest.

